# Towards Modeling in Large-Scale Genetic and Metabolic Networks

**DOI:** 10.1101/2025.11.11.687908

**Authors:** Marco Polo Castillo-Villalba

## Abstract

Mathematical modeling of large-scale genetic and metabolic networks remains an open area of research with profound implications for understanding biochemical signaling systems in both prokaryotic and eukaryotic organisms. In this work, we extend the biochemical modeling framework based on the **Power Law formalism**, originally developed by M.A. Savageau to elucidate design principles in small bacterial networks. This formalism captures gene regulation and metabolic fluxes through two fundamental sets of parameters: *kinetic parameters* and *kinetic orders*. The kinetic parameters determine the magnitudes of fluxes, while the kinetic orders appearing as exponents in the power-law representation encode the regulatory structure of the system. We propose a mathematical technique to characterize the entire family of systems associated with the space of kinetic orders. We show that this exponent space possesses a well-defined structure in **toric geometry** and can therefore be modeled using **algebraic varieties**. To represent large-scale networks, we associate a toric variety with each polynomial equation of the system and then use geometric techniques to coherently combine these varieties into a global structure, or **fan**. This construction enables a modular and geometrically consistent representation of complex biochemical networks.

## 1. Introduction

### 2. Theoretical Background: Biochemical Networks as Toric S-systems

One of the main ideas behind the analysis of biochemical systems is the conservation of mass in reactions. In addition, the Michaelis–Menten rate law is one of the fundamental concepts in biochemical systems. However, in recent decades, this law has been refined to provide more detailed biochemical information about the biomolecules participating in the network. For example, it is now possible to consider reactions with arbitrary orders, such as the number of binding sites in gene operator regions or activation parameters from Hill functions.

This is the basis of the Power Law formalism [**11**], which, through parametric extensions, allows for the simultaneous modeling of both metabolism and gene regulation. The formal expression of this model is given below:

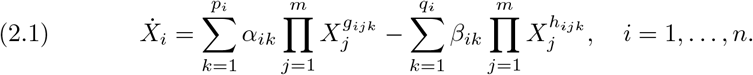

Here, the kinetic parameters *α*_*ik*_ and *β*_*ik*_ capture the rate constants for production and degradation, respectively, of each species or biomolecule in the network. The kinetic orders *g*_*ijk*_ and *h*_*ijk*_ represent genotypic information, such as binding sites and the kinetic orders of catalyzed reactions or multimeric regulation sub units in protein structures. Therefore, Equation (2.1) captures all the information required to analyze genetic–metabolic networks.

#### Definition 2.1. (S-systems).

We define the number of combinations of dominant terms in Equation (2.1) as:

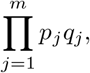

which partitions the space of concentrations (*X*_1_, …, *X*_*n*_). Each partition corresponds to a dominant term, or **S-system**, as defined in [**13**], [**11**], [**7**], [**8**], [**9**]. This is expressed as follows:

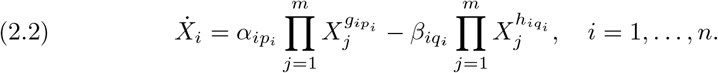

In the context of this formalism, an S-system defines a quantitative phenotype and corresponds to a biological reality in molecular biology. These systems are not merely mathematical artifacts; they represent rates of change in molecular concentrations derived from this formal gradient.

#### 2.1. Toric S-systems

In this section, we highlight the importance of studying the exponents 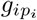 and 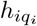 in an S-system. We assert that the monomial map 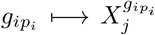 represents a **Genotype–Phenotype** relationship with properties of great interest for further study. This central idea forms the foundation of our research. Below, we briefly summarize the definitions introduced in well-known works such as [**7**], [**13**], [**11**], and [**7**].

##### Definition 2.2. (Exponent Space or Support)

For an S-system defined by 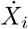, we define the *exponent space* of 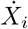, also formally known as the *support* of a polynomial, as:

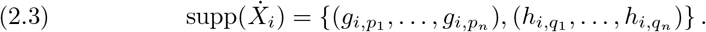

Using the support of any S-system supp 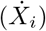, we can construct all linear combinations from the two vectors in the support, 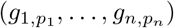 and 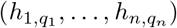. The set of all such vector combinations defines the **cone** associated with 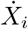, as described in [**4**].

##### Definition 2.3. (Cone)

Let 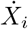 be an S-system as defined above. We define the cone associated with it as:

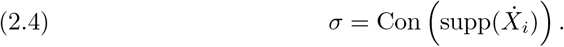

This cone represents the linear space generated by all combinations of the vectors in the support space, including addition, subtraction, and scalar multiplication. If the vectors consist only of integer entries, the cone is called a *lattice cone* [**4**].

##### Definition 2.4. (Dual Cone)

Let *σ* ⊆ℤ^*n*^ be a lattice cone. The dual cone *σ*^∨^ is defined as:

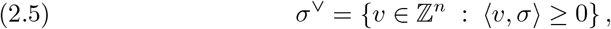

as described in [**4**].

##### Definition 2.5. (Monoid)

By intersecting the dual cone with the *n*-dimensional integer lattice, we obtain:

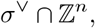

which we call a **monoid**. This monoid, denoted by *M*, has rank *n* and is isomorphic to ℤ^*n*^ [**4**].

##### Lemma (Gordan’s Lemma and the Hilbert Basis)

Given a cone *σ*, the intersection of its dual with the *n*-dimensional integer space,

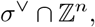

forms a monoid that has a finite set of generators. This generating set is called the **Hilbert basis** [**17**], [**4**].

###### Definition 2.6. (Fan or topological environment)

Given a collection of dual cones

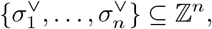

we define the *fan* 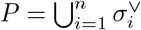 as the union of all these cones. We denote this fan by Σ ⊆ ℤ^*n*^ [**4**].

Finally, we construct a coordinate polynomial ring for S-systems. Let 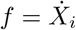 be an S-system defined for any *i*-th species in a metabolic network. Given the support of this polynomial supp(*f*) and its associated cone *σ*, we define the polynomial ring:

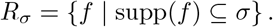

In other words, the exponents of an S-system (i.e., the kinetic orders) are embedded in a geometric space. We will explore new topological properties that arise from this geometric object in the following sections.

###### Definition 2.7. (Toric Ideal).

A toric ideal is the set of prime binomials defined by *I* =*< X*^*A*^ − *X*^*B*^ |*H* ∗ (*A* − *B*) = 0 *>*, where *H* is the *Hilbert basis* of the polynomials generated from *I*, [**17**], [**3**].

###### Definition 2.8. (Toric Variety).

We define a **toric variety** associated with a polynomial *f* as follows:

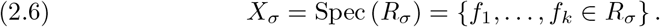

Here, the notation Spec (*R*_*σ*_) denotes the set of all prime polynomials (i.e., non-divisible polynomials) {*f*_1_, …, *f*_*k*_} in the ring *R*_*σ*_ for some integer *k*, see [**4**].

###### Definition 2.9. (Environment Cone).

Let *f* be an S-system representing the rate law for the change in concentration of any species in a gene–metabolic network. We compute the support of *f* and the cone associated with it, *σ* ⊆ℤ^*n*^. Since supp(*f*) ⊆*σ*, we define the **environment cone** associated with *f* as the dual cone *σ*^∨^ of *σ*. Thus, we have the inclusion relation:

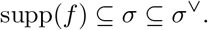

##### Relevance in Enzyme Kinetics

Up to this point, we have equipped the set of vectors associated with the kinetic orders of an S-system with new geometric properties. We have defined cones, polynomial rings, and toric varieties. The most important result arises from **Gordan’s Lemma** [**4**], which guaranties the existence of a generator basis (the **Hilbert basis**) for the entire kinetic order space defined for any species in the network. Next, we will explore how to exploit the properties induced by this Hilbert basis.

##### Toric S-systems and Dominant Fluxes

Given the Hilbert basis *H* = {*h*_1_, …, *h*_*k*_}, we represent it in matrix form considering each *h*_*i*_ as a row vector:

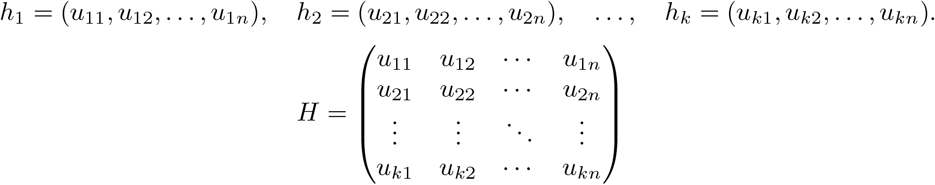

We can use this matrix to represent the variables *X*_*i*_ in a new coordinate system using the following monomial transformation:

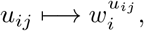

as discussed in [**4**], [**17**], and [**3**]. This transformation reveals dynamical properties encoded within the S-systems through the Hilbert basis [**17**], [**5**].

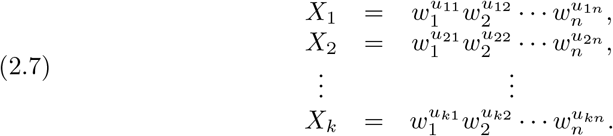

This matrix representation allows embedding all S-system equations into a new system of monomial coordinates. Hence, we extend the analysis of molecular networks toward large-scale genetic and metabolic systems.

###### Theorem 2.10. (Toric S-system).

*If we substitute the above change of coordinates into Equation (2.2), the original S-system equations, we obtain the following expression, valid for all i* = 1, …, *n:*

***Proof***. *Given an S-system representation of molecular networks as in Equation (2.2), and applying Hironaka’s theorem together with the algorithmic construction of the Hilbert basis (a finite number of times) to the S-system equations* [**1**], [**6**], [**5**], *we find that every S-system is toric and can be described by:*

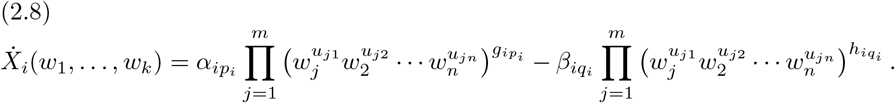

*Or, more compactl:*

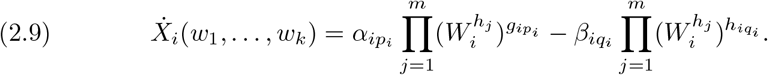

*We thus define a* ***Toric S-system*** *according to the expression above, where h*_*j*_ *is an element of the Hilbert basis for some j. We assert that the binomials constructed from the Hilbert basis H*,

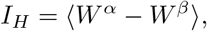

*with* 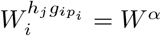 *and* 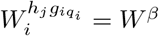, *are* ***toric binomials. q.e.d***.

The monomial transformation,

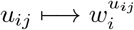

represents, in this geometric framework, the **Genotype–Phenotype relationship**. These are the coordinate transformations of the geometric basis corresponding to the kinetic orders such as the number of binding sites in regulatory genomic sequences, the order of biochemical reactions in metabolism, or the degree of multimeric regulation in proteins.

The monomials 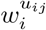 represent the chemical concentrations of the species or biomolecules described by the S-system *X*_*i*_. The coordinates *u*_*ij*_ encode information derived from the genome sequence, which is then expressed through the phenotypic gradient:

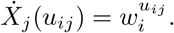

Thus, the mapping of dynamical behavior occurs through this gradient, linking genotypic parameters to phenotypic observables.

## 3. Discussion

We have presented new topological properties of biochemical networks by exploiting the structural features of the Power Law and S-system modeling frameworks. In particular, we emphasize the significance of the monomial map: 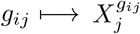, which provides a bridge between biochemical kinetics and algebraic geometry. By focusing our analysis on the exponent space of S-systems, we study the kinetic orders using computational algebraic geometry techniques. This allows us to identify geometric bases that constrain and characterize the possible values of these parameters, thereby facilitating the solution of large systems of differential equations.

The central open questions that can be addressed with this approach are as follows:

- Under what conditions are these geometrical regions valid for assigning values to the kinetic orders?
- Which of these regions represent true biological information and which are merely mathematical artifacts?

Another advantage of the proposed methodology is its potential to model the dynamics of large-scale metabolic and gene-regulatory networks. The monomial map has a formal justification through the mathematical concept of a **toric variety**. Although toric varieties are abstract algebraic objects, they serve as powerful tools for modular analysis: each variety corresponds to a submodule of the biochemical network, and the full network can be reconstructed by *gluing together* these varieties using geometric techniques.

While a single toric variety may provide limited insight into the overall biochemical system, the ensemble of all varieties properly joined captures the global dynamics of large-scale genetic and metabolic networks. This modular, geometrically grounded framework thus offers a new perspective for understanding and simulating the complex interplay between genotype and phenotype.

Finally, we have included in the Supplementary Information the complete set of equations used for the toric analysis of a gene–metabolic network in the allosteric system of the tetrameric protein TrpR (tryptophan repressor) in E. coli K-12. In a future work, we will solve the complete dynamics using toric varieties such as biochemical modules.

## Supporting information

Latex_Manuscript

## 4. Acknowledgment

Marco Polo Castillo Villalba is a doctoral student from the Programa de Doctorado en Ciencias Biomédicas, Universidad Nacional Autónoma de México (UNAM) and has received the CONAHCYT fellowship 754050. We acknowledge funding from UNAM and from the National Institutes of Health (grant number 5R01GM110597-03). I give a special thanks to Michael A. Savageau for all technical comments during my scholar stay with him, Pedro Miramontes Vidal and Julio Collado-Vides and technical contribution by Laura Gómez Romero.

## Support information

**Figure 1.**
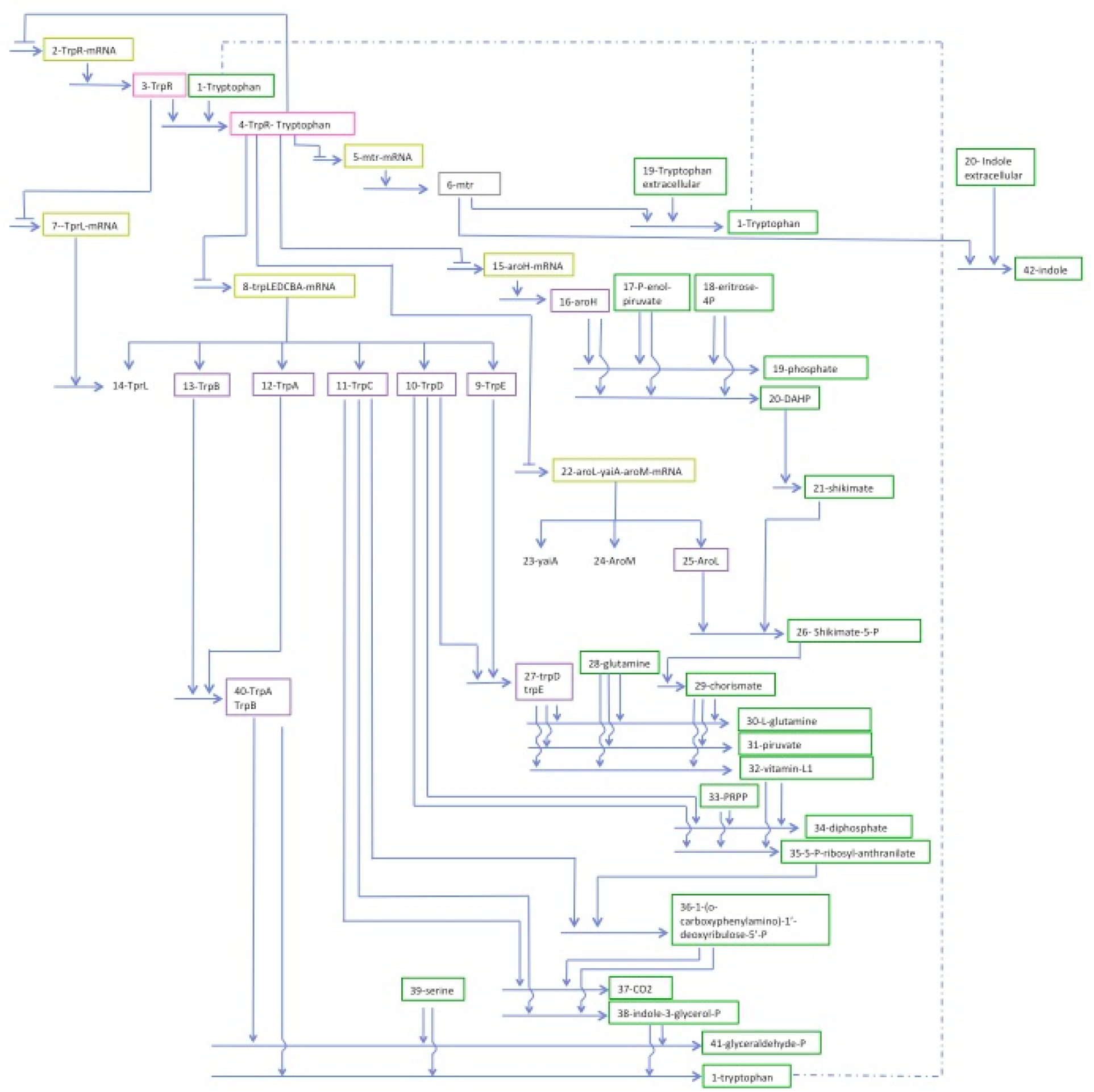
Tryptophan genetic and metabolic network.

**Figure 2.**
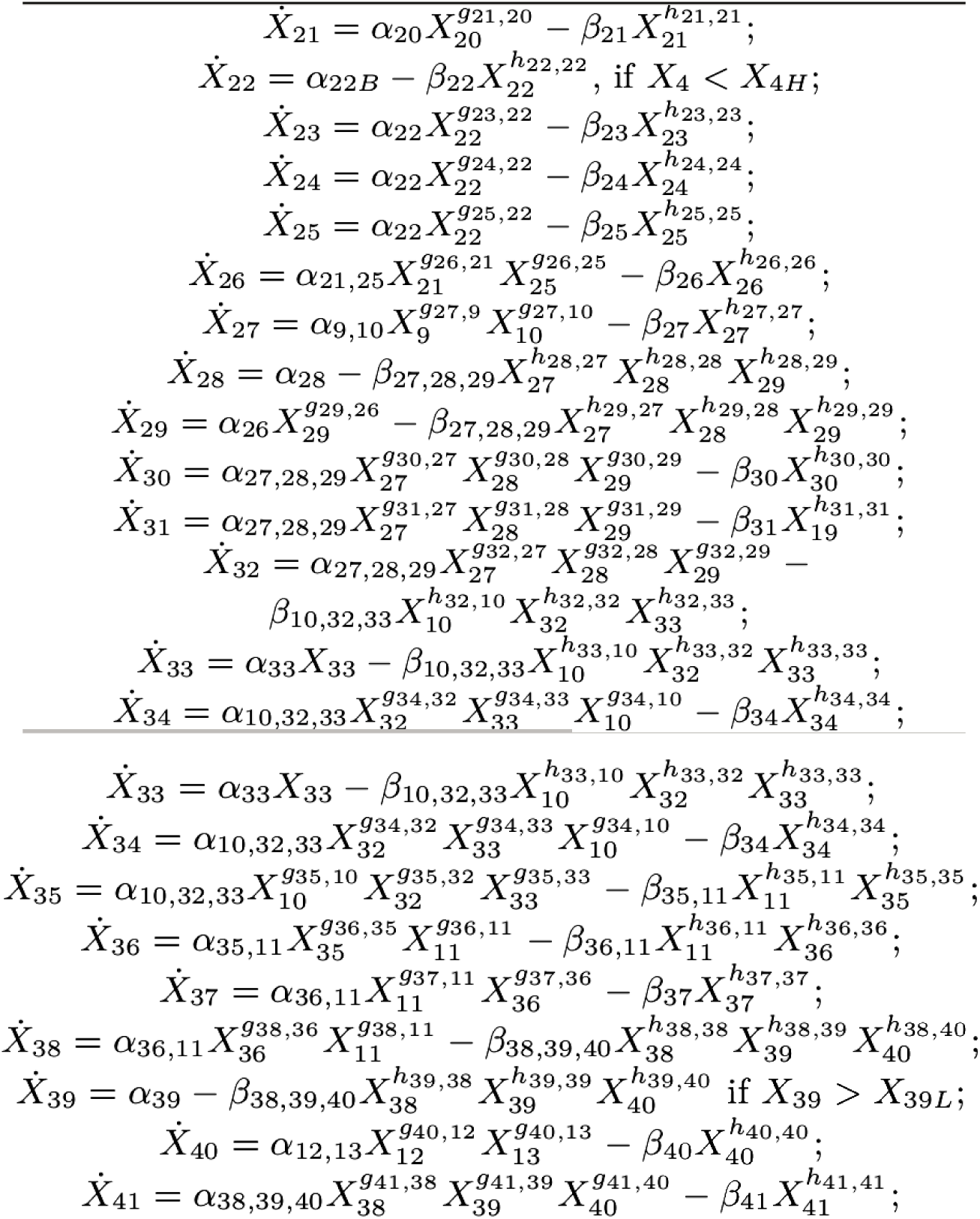
Toric S-system equations for tetrameric tryptophan protein.

