## Supplementary material for "Towards Modeling in Large-Scale Genetic and Metabolic Networks": Latex_Manuscript: Topological_Env_kineticorders.pdf

Marco Castillo-Villalba

Abstract. We summary the results of this work, as a cue to solving the lacking of kinetic orders in large-scale genetic-metabolic networks from eucaryotes and prokaryotes and to be available to model quantitative phenotypes, we used the formalism of previously works mentioned here. As a corner stone of our project, it is sustained in the application and theoretical approach from S-systems formalism, to solve dynamics on metabolic networks. In recent years have been developed a software to solve dynamics for S-systems (Design Space Toolbox v.3.0) by M.A. Savageau [**15**], getting precise approximations of kinetic parameters (rates of production and degradation of biomolecules) for small networks, incluiding the construction of quantitative phenotypes and theirs dynamics, but is mandatory to provide the set of kinetic orders (activation parameters, reaction orders, number of binding sites in operons, etc..) for a whole solution. For the case of large-scale networks is even unsolved tha lacking of that kinetic orders. We propouse in this work a geometrical technique based in a monomial map into the concepts in topological environment, [], we postule that this map can characterize well the relationship genotype-phentoype, as consequence of this map we analyse the fix points on torus for biochemical networks, [], we conclude that these points are an equivalent of classical steady-state for phenotypes, [], it is a particular steady-state for genotypes, in this genotypic state, we did an approximation for kinetic orders, and we do an interpratation from point of view of the Bifurcation theory and molecular biology,[], [].

### 1. Introduction

We begin doing alussion to the main works developed up to now in the field of gene and metabolic networks, with that, we try to understand the left holes in the lacking of knowledge or information in the area, in the past decades. Therefore is plausible to anunciate the great contributions made by some authors, and the posibilities to do contributions in unsolved issues, exploring new properties in gene networks by means of contemporany mathematical tool, as is the case of computational algebraic geometry, we analyse a new geometrical technique to approximation for kinetic orders, making a complementary treatment with one of the authors mentioned below.

2010 *Mathematics Subject Classification.* Primary .

*Key words and phrases.* Topological Environment, S-systems, fix points on torus, quantitative phenotype; bifurcation branches; kinetic orders.

Firstly, we remember the contribution given by Rhenés Thomas and his invaluable work, **Biological Feedback**, [], since that he analized for first time gene circuits in the formalism of boolean algebra, discovering the positive regulation loops are as relevant as negative regulation loops, an example of it, is the case of activation by cAMP in genes, [], in addition the boolean schema in gene networks lets to analyse different aspects in steady-state, equilibrium points, attractors, cycles of regulations, prediction of phenotypes before mutations or knock out of genes. But mostly important is the introduction of Hill functions in the boolean analysis, letting to analyse the transcription of genes through sigmoid and S-shape curves, these functions are depending of parameter of activation $n$ in genes, it which can be related to the number of binding sites in the promoter regions in genes. The disadvantages in this theory is not consider the conduction of biomass or metabolism in networks and therefore the balance of biomass or kinetic enzyme is not taken in account.

Secondly, the analysis given by Barabasi and Kauffmann, [], [], provides evidence for the behaviour in large-scale networks, by means of concepts of random networks was understanding a hierarchical relationship between sub-groups of nodes in a called power law, said in another way, the subgroups are governed in clusters, separated in a rational distance $\gamma$ or with fractional dimension, this number is depending of a constant parameter, named degree of conectivity in the network $c$, in this approach the models of diffusion and thermodynamics have been studied, but the computational complexity in them, in several cases makes intractable. However in the study of diffusion of cells of modelling highest levels of organization have been usefull, [].

In the analysis of networks with fluxes of biomass, the theory of Flux Balance Analysis, developed by B.O. Palsson and his team, has provided important results in metabolism modelling, [], [], but mainly in the development for the synthetic biology, these results are direct consequence of the formalism in which this methodology is sustained, the theory of optimization in networks is very successfully by tradition in the design of optimal routes, minimal and maximal subnetworks for supply chains, as consequence is viable as tool for bio-engeenering, but nowadays is complicated the tratment of gene regulation in this scope.

For the case of analysis in gene-metabolic networks, the formalism of power law and S-systems has gone developing during several years, since 1976 by M.A. Savageau, [], [], the amount of articles and examples studied in microbiology has been too many, the inclusion of gene an metabolism in the mathematical modelling lets a wide range of analysis in diverses example of interst in biochemical systems, the background of this theory is based in fundamental priciples of enzyme kinetics and multivarite calculus, in addition with experimental evidence of their results and curation of data from biomedical literature. It makes a powerfull tool for the understanding of prinples of design in gene regulation, thus as the development of ideas for new cellular theories. The limitations of this focus are several, since that the great amount of kinetic parameters and orders increase fastly in proportion with the size of the network. In recent years, M.A. and his team has developed a software, Design space toolbox, v 2.0, [], for the analysis of small networks, solving the complete dynamics and elucidating quantitative phenotypes of the network, at same time this programm can approximate in numerical ranges, the complete set of kinetic parameters (production and degradations rates in species). However the inclusion of kinetic orders (number of binding sites, number of orders of reaction, parameters of activation) are obtained in an idependent way in the model, this fact is a limitation for the analysis in large-scale networks. We propose here, a complentary technique for the analysis of networks with Design space, exploting new topological properties in the network by means of concepts of Topological environment from S-systems, developed in supplementary material,[]. We believe that the methodolgy exposed here in addition the techniques developed by S-systems and implemented on Design-space platform, we can scale the simulations of dynamics for large-scale networks in mechanisist approach as is the case of pure biochemical reactions.

On the development of this article we do mention to three previously works, "Genetic-Metaolic Networks can be modeled as toric-varieties", [], "Stochastic Environmnet Matrices for Genetic-Metabolic Networks ", [], and "Topological Environment of genetic-metabolic networks", [], published on the Arxiv and Bio-Arxiv platforms. With the theoretical formalism explained in thoses works, we are able with the sufficient tools to analyse a study case for a small genetic-metabolic circuit, it which will be very usefull to valid bounds for kientic orders and to elucidate the reaches of this mathematical modeling in gene networks. In the study case exposed here, the variables of the model has been well biologically justified by authors as, M.A. Savageau , [], [], such as the dynamics and the experimental results. Theses variables mainly are: kinetic paramters and kinetic orders associated to the expression of two regulators gene.
With the methodology given here, we have as main aim to give a technique to bound the values of kinetic orders in a specific invariant geometrical regions, as consequence of new topological properties from network, it is the case of fix points on torus, [].

In the Section 2.0, we start a brief summary of the theoretical background in the scope of the Power Law Formalism and S-systems, followed of the main concepts in toric algebraic geometry in S-systems (Toric S-systems), we also do mention, about the possible relationships there is exists between, bifurcation theory, [], and catastrophe geometries, [], with all those concepts, at the same time we give a biophysics interpretation.
On these tools at hand, we prepare the landscape for the Section 3.0, to describe an algorithmic approach to build geometrical regions where the set of kinetic orders can take numerical values, therefore they can be used for a specific S-system associated to a biomoelcule in the network, in this part, in the sub-section 4.1 we decribed a example of application, a small gene circuit well known in the gene regulation research, it which consist of two regulators, the rate of expression of these genes is modeled through S-systems equations, in the subsections we exhaustively computed a set of stochastic biochemical environment matrices (environment dual cones) associated to the S-systems of the example with a normal distribution, we provide a set of 20 thousands matrices for every equation that describes each regulator, we describe the constrcution of environment matrices with detail to any

specie in a genetic-metabolic network.

Once computed the stochastic envrionment matrices of our example, we are ready in the Subsection 4.3 to analyse a particular invariant geometrical regions, in this biochemical environment we proved the existence of a set of geometrical basis called Hilbert basis, [], they are computed on the Singular computer platform (a suit in computational algebraic geometric algoriothms) [], on these Hilbert basis, there is exist a minimal unimodular sub-matrix (with determinant $\pm 1$), this set is contained in the most of all the set of Hilbert basis associated to the 20 thousands stochastic matrices computed in the sub-section 4.2. We shown that this unimodular sub-matrix is a small geometrical cluster of vectors, they contains numerical values for the kinetic orders in our example, we will notice said values are in agreement with the biomedical literature, [], [], and gene regulation process defined by the network of our example.

In the next sub-section, 4.4, we use the Design space toolbox v.4. software, [], to analyse the dynamics of our study case, we take a range of different values computed in the sub-section 4.3 for kinetic orders. Hence, with all this biochemical information, we elucidated the whole set of quantitative phenotypes yield on this gene circuit. The phenotypes are produced in accoording to the rates of production and degradation for the expression of the two genes in our example in the beginning of the section.

In the last part, we discuss about of the reaches of the methodology in the solution for large-scale networks and theirs dynamics. So we also consider the possibility to elucidate when the minimal geometrical regions might be mathematical artifacts.

### 2. Theoretical background of Biochemical networks and Toric S-systems.

One of the main ideas behind analysis of biochemical system is the balance of mass in the reactions, in additon Micheles-Menten rate law, is an of the concept fundamentals in biochemical systems, but in the last decades this law has been improved to give more biochemical details of the biomolecule participatiyng in the network, for example now we can consider any arbitrary order of reaction, as to consider the number of binding sites of operator region in genes, or an activation parameter of Hill's function. It is the case of the theory of power law formalism, [], where considering this parametric extension, in this modelling is possible to include matabolism and gene regulation togheter. The formal expression of this model is presented below.

$$\dot{X}_i = \sum_{k=1}^{p_i} \alpha_{ik} \prod_{j=1}^{m} X_j^{g_{ijk}} - \sum_{k=1}^{q_i} \beta_{ik} \prod_{j=1}^{m} X_j^{h_{ijk}},\ i = 1, ..., n. \tag{2.1}$$

Here, the kinetic parameters, $\alpha_{ik}$, $\beta_{ik}$, capture information of constant rate laws of production and degradation repectevely, of any species or biomolecule in the network, the kinetic orders $g_{ijk}$, $h_{ijk}$, capture genotypic information, from bindings sites and kinetic orders of catalysed reactions, as we have commented at first. Hence, the expression 2.1, has captured all information to analyse genetic-metabolic networks.

Definition 2.1. (**S-systems**). We define a number of combinations of dominant terms in 2.1, given by:

$$\prod_{j=1}^{m} p_j q_j;$$

partitions of the space of concentrations $(X_1, ..., X_n)$, to each partition we call it, a dominant term or **S-system**, [**15**], [**14**], [**9**], [**10**], [**11**]; as shown below,

$$\dot{X}_i = \alpha_{ip_i} \prod_{j=1}^{m} X_j^{g_{ip_i}} - \beta_{iq_i} \prod_{j=1}^{m} X_j^{h_{iq_i}},\ i = 1, ..., n. \tag{2.2}$$

In the context of this formalism, an S-system defines a quantitative phenotype, and also corresponds to a reality of the molecular biology, not only as mathematical artifacts, clearly they are rate of molecular concentrations computed by this gradient.

**2.1. Toric S-systems and Bifurcation theory : A biophysical hypotesis.** In this section we remark the importance to study the of the exponents $g_{ip_i}$ and $h_{iq_i}$ in an S-system, and we do the assertion that the monomial map $g_{ip_i} \longmapsto X_j^{g_{ip_i}}$ is a relationship **Genotype-Phenotye** with interesting properties to study, and this central idea, is the fundamental ground of our research. We summary in brief definitions given in past works, [], [].

Definition 2.2. **(Exponent-space or support)** For an S-system given $\dot{X}_i$, we define the exponent-space of $\dot{X}_i$ or formally called support of a polynomial, as,

$$supp(\dot{X}_i) = \{(g_{i,p_1}, ..., g_{i,p_n}), (h_{i,q_1}, ..., h_{i,q_n})\} \tag{2.3}$$

With the support of the any S-system $supp(\dot{X}_i)$, we can build all the linear combinations from the two vectors in the support, $(g_{1,p_1}, ..., g_{n,p_n})$ and $(h_{1,q_1,...,h_{n,q_n}})$, this set of combinations of vectors define the **Cone** associated to $\dot{X}_i$.

Definition 2.3. **(Cone)**. Let $\dot{X}_i$ be an S-system defined as above, we define the cone associated to it, as follows,

$$\sigma = Con\left(supp(\dot{X}_i)\right). \tag{2.4}$$

It is the linear space defined by all combinations for the vectors of support space, incluiding addition, substraction and scalar multiplycation of vectors, if the vectors consist only of integer entries, is called a lattice-cone.
Now we intersect the cone with the $n-$dimensional integer space, we write, $\sigma \cap \mathbb{Z}^n$, we call to this set as a monoid.

We build a coordinate polynomial ring of S-systems. Let be $f = \dot{X}_i$ an S-system defined for any $i-$esim specie in the metabolic network, given the support of this polynomial $supp(f)$ and the cone associated to this set, $\sigma$, we define a polynomial ring, as follows, $R_\sigma = \{f \mid supp(f) \subseteq \sigma\}$. In other words, the exponents of an S-system or kinetic orders are embedding in a geometrical space, we will exploit new topological properties on this new geometrical object, as we can see.

Definition 2.4. **(Toric variety in S-systems)**. , we define a **toric variety** associted to $f$ as follows,

$$X_\sigma = Spec\,(R_\sigma) = \{f_1, ...., f_k \in R_\sigma\}. \tag{2.5}$$

where the notation for $Spec\,(R_\sigma)$ means, the set of all the prime polynomials (non-divisible polynomials) $\{f_1, ...., f_k\}$ in the ring $R_\sigma$ for some integer $k$.

Definition 2.5. **(Dual cone)**.

$$\sigma^\vee = \{v : < v, \sigma > \quad >= \quad 0\}. \tag{2.6}$$

Definition 2.6. **Environment cone.** Let $f$ be a S-system of any rate law of a chemical concentration change of any species in a gene-metabolic network. We compute the support of $f$ and the cone associated with it, $\sigma \subseteq \mathbb{Z}^n$. We know that $supp(f) \subseteq \sigma$. We define an **environment cone** associated to $f$ as the dual-cone $\sigma^\vee$ of $\sigma$, we also know that $supp(f) \subseteq \sigma \subseteq \sigma^\vee$.

**Hilbert basis** (Gordan's Lemma). Given a cone $\sigma$ and we take the intersection with the $n-$dimensional integer space $\sigma \cap \mathbb{Z}^n$, then this monoid, has a finite set of generators, this set of generators is called, the **Hilbert basis** .

**Relevance in kinetic enzyme.** Up to here, we have equipped of new geometrical properties to the set of vectors associated to kinetic orders for an S-system, we have defined, cones, ring of polynomials, and a toric variety. But the most importance is the fact through of Gordan's lemma we have found a basis o generators (Hilbert basis) for all space of kinetic orders defined for any specie on the network. Now, we will try to exploit the properties gifted by the Hilbert basis.

**Fix points on torus**. There exists several programms and algorithms to compute Hilbert basis, [], once obtained the set of generators $H = \{h_1, ..., h_k\}$ for some cone $\sigma = Con(b_1, ..., b_r)$ where each vector $b_j$ is a face, [], that generates the cone. we define the fix points on the torus or toric variety through of the following formula, see [].

$$X_{b_j} = \lim_{Z \longrightarrow 0} Z^{<b_j, h_i>} = (\lim_{Z \longrightarrow 0} Z^{<b_j, h_1>}, ..., \lim_{Z \longrightarrow 0} Z^{<b_j, h_k>}). \tag{2.7}$$

We compute this limit for each face $b_j$ in the cone, therefore, there are $r$ fix points, or in another way, $X_{b_1}, ..., X_{b_r}$. These limtis exists, if and only if, the set of constraints fullfil the conditions as we can see below.

$$< b_j, h_1 >>= 0, < b_j, h_2 >>= 0, ..., < b_j, h_k >>= 0. \tag{2.8}$$

For some integers, $j$ and $k$, and usual scalar inner product, $<,>$. We remember that the vectors $b_j$ are of fact the **kinetic orders** associated to the support of some S-system $f$ as we saw above. We use this set of constraints in addition with the constraints for dominant fluxes, to determine values for kinetic orders in the algorithm described in this work.

**Assertion.** The classical point fixs, are information characterized by the well known steady-state from phenotype, in our case the fix points on torus, are information provided from genotype, we mean, this is another consequence of the monomial map, $g_{ip_i} \longmapsto X_j^{g_{ip_i}}$ studied here.

**Toric S-systems and dominant fluxes**.

With the Hilbert basis at hand, $H = \{h_1, ..., h_k\}$, we give a matricial representation of it, if we consider each $h_i$ as a row vector, thus we write, $h_1 = (u_{11}, u_{12}, ..., u_{1n})$, $h_2 = (u_{21}, u_{22}, ..., u_{2n})$,...,$h_k = (u_{k1}, u_{k2}, ..., u_{kn})$.

$$H = \begin{pmatrix} u_{11} & u_{12} & ... & u_{1n} \\ u_{21} & u_{22} & ... & u_{2n} \\ ... & ... & ... & ... \\ u_{k1} & u_{k2} & ... & u_{kn} \end{pmatrix}$$

To continuation, we can use the matrix defined above to represent the varibales $X_i$ in a new system of coordinates, by means of the following monomial transformation, $u_{ij} \longmapsto w_i^{u_{ij}}$, see [], the relevance to do that, is to notice dynamics properties encrypted for S-systems into of the Hilbert basis, we will see in the last part of this section.

$$\begin{array}{llll} X_1 = w_1^{u_{11}} * & w_2^{u_{12}} * & ... & * w_n^{u_{1n}}, \\ X_2 = w_1^{u_{21}} * & w_2^{u_{21}} * & ... & * w_n^{u_{2n}} \\ ... & ... & ... & ... \\ X_k = w_1^{u_{k1}} * & w_2^{u_{k2}} * & ... & * w_n^{u_{kn}} \end{array} \tag{2.9}$$

Definition 2.7. **(Toric S-system)**. If we subtitute this change of coordinates in the expression 2.2, the original S-systems, we have the following equation, it which is valid for all $i = 1, .., n$,

(2.10)

$$\dot{X}_i(w_1, ..., w_k) = \alpha_{ip_i} \prod_{j=1}^{m} (w_j^{u_{j1}} * w_2^{u_{j2}} * ... * w_n^{u_{jn}})^{g_{ip_i}} - \beta_{iq_i} \prod_{j=1}^{m} (w_j^{u_{j1}} * w_2^{u_{j2}} * ... * w_n^{u_{jn}})^{h_{iq_i}}.$$

Or simply,

$$\dot{X}_i(w_1, ..., w_k) = \alpha_{ip_i} \prod_{j=1}^{m} (W_i^{h_j})^{g_{ip_i}} - \beta_{iq_i} \prod_{j=1}^{m} (W_i^{h_j})^{h_{iq_i}}. \tag{2.11}$$

We define a Toric S-system, to the expression given above, where the $h_i$ is an element of the Hilbert basis, and we used the notation $W_i = w_1^{u_{i1}} * w_2^{u_{i2}} * ... * w_n^{u_{in}}$.

Definition 2.8. **(Bifurcation.)**. Let $G$ be a Lie group acting on a vector space $V$. A **bifurcation** problem with symmetry group $G$ is a an abelian group of functions $X \in \Lambda_{x,\lambda}(G)$ satisfying $X(0,0) = 0$ and $(dX)_{0,0} = 0$, see details [].

We do the assertion that any toric S-systems satisfies this definition, and we explain more facts in the following, it which shows this result and its implications in gene regulation.

**Action on algebraic torus and orbits.** The products in the exponents of the last expression in 2.11, $h_i * g_{ip_i}$ and $h_i * g_{ip_i}$, are called an action of group on torus, and they define orbits in the complex $n-$dimensional projective space, for technical details, see []. This fact is a result whose implications are of interest for us, as the systems are highly non-linear polynomials, is hard to understand their equilibrium points and singularities, the algebraic torus is a **compact Lie group**, it implies, that we can solve the kind of bifurcations in these family of polynomials. The study of branches of bifurcation in S-systems, is a vital key for the understanding of the evolution of dynamics in gene networks.

The branches of bifurcations are the main directions to conduct the dynamics, agreement to the Arnold's Classification (Catastrophe theory), [], he defined a set of **Catatrophe geometries**, $A_0, A_1, A_2, A_3, A_4, A_5, A_K, D_4^-, D_4^+, D_5, D_K, E_6, E_8$, We do the assertion about the set of family of S-systems belongs to the $E_8$ catastrophe, $E_8$ are geometries associated to compact Lie groups, consequence from that the S-systems has been embedding in the algebraic torus.

We consider the following biochemical flux, given by, $\dot{X}(x,y,z) = z^5 + x^{15} + xy^7 - y^6 z$, we plotted in the fig 1, the flux in 3d dimensional Euclidian-space, in the fig 2, its projection in the 2d dimensional space, in the third one figure, we plotted the bifurcation branches associated to the flux $\dot{X}$, as we can see, the branches in fig. 3 conductes the dynamics in the plot 3d-dimensional.

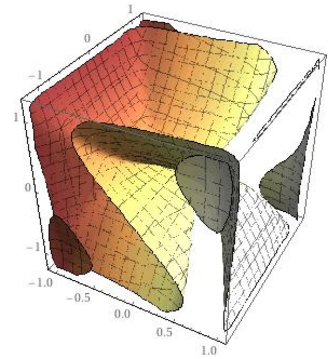


Figure 1. Biochemical flux from a power law equation representation, $\dot{X}(x,y,z) = z^5 + x^{15} + xy^7 - y^6 z$, with kinetic parameters all with value equal to 1.

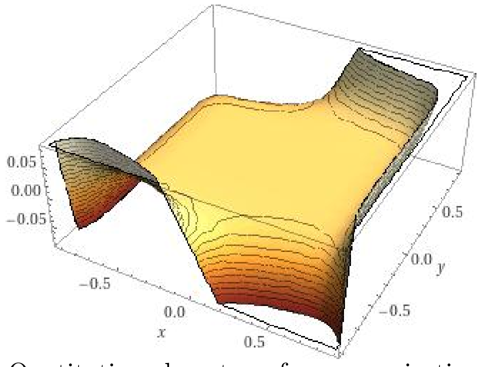


FIGURE 2. Quatitative phenotype from a projection S-system of 3d dimensional flux in the fig. 1.

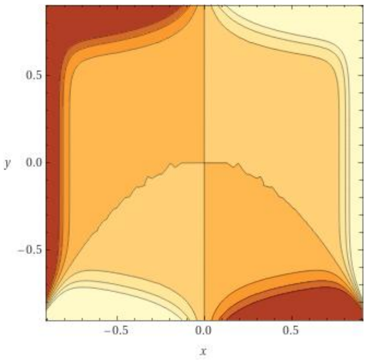


FIGURE 3. Branches of bifurcation projected in the 2d real-plane, mapped from fig. 2. The kinetic orders computed from Hilbert Basis determiens these bifurcation branches.

**Biophysical interpretation**.

**Hypotesis 1.** The rate of initial interaction of transcripts with the region of binding sites or domains in genes and is related to the minimun energy of interaction, it determines the kind of bifurcations in S-systems and therefore the directions in the dynamics.

**Hypotesis 2.** The group of branches of bifurcation for high dimensional S-systems is computed by a minimal region for kinetic orders and they are bounded by the Hilbert basis, that group of branches determines the dynamics in S-systems.

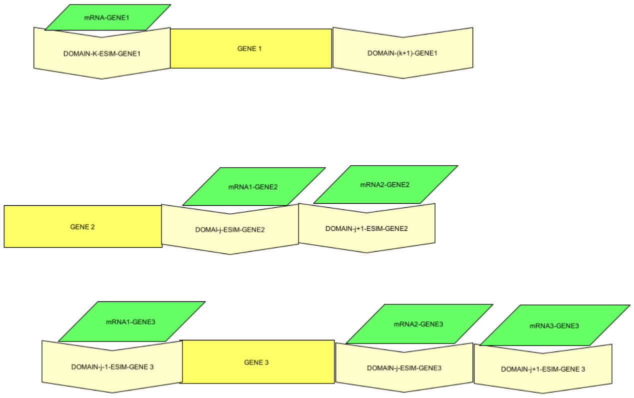


FIGURE 4. Different modos of initial interaction in the gene transcription implies and determines different bifuraction branches in S-sytems, they are produced by the minimal values for kinetic orders, bounded by gemetrical regions.

For example in the figure 4, we have threes simply sceneries, where the S-systems associated, are: $\dot{X}(x) = \alpha * x - \beta$, $\dot{X}(x) = \alpha * x^2 - \beta$, $\dot{X}(x) = \alpha * x^3 - \beta$, with set of kinetic orders given by, $\{g_1, g_2, g_3\} = \{1, 2, 3\}$, the kinetic orders, let us to know the geometry of the branches associated, in this case: the constant line $\alpha$, straight line with slope $2 * \alpha$, and a parabola with branches $\pm\sqrt{3} * \alpha$, they coincide with the directrixes.

**Assertion.** The Hilbert basis and the fix points on torus, we can declare that they are topological **motifs** in the network, and they are invariants under the biochemical environment of the biomolecule considered in the genetic-metabolic network, we made thousands of simulations presented through of this work to have evidence on that, but mathematical theorems confirm this fact.

The monomial transformation $u_{ij} \longmapsto w_i^{u_{ij}}$ represents in this geometry, the relatioship **Genotype-Phenotype**, they are coordinates of geometrical basis for the kinetic orders, the monomials $w_i^{u_{ij}}$ are chemical concentrations of any species or biomolecules described by the S-system $X_i$, the coordinates $u_{ij}$ are information from genome sequence it which is read by the phenotypic gradient $\dot{X}_j(u_{ij}) = w_i^{u_{ij}}$, as we saw in the different scenaries in the fig. 4, the dynamcis are mapped by this gradient.

**Dominant Fluxes in toric S-systems.** Now we consider the equation (2.1), and we use the monomial transformation (2.9), we substitute the new coordinates system in (2.1),

$$\dot{X}_i = \sum_{k=1}^{p_i} \alpha_{ik} \prod_{j=1}^{m} (W_i^{h_j})^{g_{ijk}} - \sum_{k=1}^{q_i} \beta_{ik} \prod_{j=1}^{m} (W_i^{h_j})^{h_{ijk}},\ i = 1, ..., n. \tag{2.12}$$

We run some terms on the sum, thus we have the following formal expression,

$$\dot{X}_i = \alpha_{i1} \prod_{j=1}^{m} (W_i^{h_j})^{g_{ij1}} + \alpha_{i2} \prod_{j=1}^{m} (W_i^{h_j})^{g_{ij2}} + \alpha_{i3} \prod_{j=1}^{m} (W_i^{h_j})^{g_{ij3}} + .... -$$

$\beta_{i1} \prod_{j=1}^{m} (W_i^{h_j})^{h_{ij1}} - \beta_{i2} \prod_{j=1}^{m} (W_i^{h_j})^{h_{ij2}} - \beta_{i3} \prod_{j=1}^{m} (W_i^{h_j})^{h_{ij3}} - .....,$ (2.13)

We get several inner products in the exponents of this expression, as we show below,

$$\langle h_j, g_{ij1} \rangle,\ \langle h_j, g_{ij2} \rangle,\ \langle h_j, g_{ij3} \rangle \ ...,\ \langle h_j, h_{ij1} \rangle,\ \langle h_j, h_{ij2} \rangle,\ \langle h_j, h_{ij3} \rangle \ .....$$

In the following we take the minimals of all this set of scalar products, to construct branches of toric S-systems, we call to that branches **dominant fluxes**, as we can see in the next example,

Given the expression for the biochemical flow in a biomolecule $X_3$, we compute the Hilbert basis associated,

$$\dot{X}_3(u_1, u_2, u_3, u_4) = c_1 * u_6 + c_2 * (u_2^2 u_3^2 u_4^2 u_6^2) u_6 + c_3 * u_4^2 u_6 - c_4 * u_3 u_4^2 u_6. \tag{2.14}$$

$$[\mathrm{H}_{\dot{X}_3}]_{8\times 8} = \begin{pmatrix} 1 & 0 & -1 & 0 & 0 & 0 & 0 & -1 \\ 0 & -1 & 0 & 0 & 0 & 0 & 0 & 0 \\ 0 & 0 & 1 & 0 & 0 & 0 & 0 & 0 \\ 1 & 0 & 0 & -1 & 0 & 0 & 0 & 0 \\ 1 & 0 & 0 & 0 & -1 & 0 & 0 & 0 \\ 1 & 0 & 0 & 0 & 0 & -1 & 0 & 0 \\ 1 & 0 & 0 & 0 & 0 & 0 & -1 & 0 \\ 0 & 0 & 0 & 0 & 0 & 0 & 0 & 1 \end{pmatrix}$$

(2.15)

We define the dominant branch or main branch for $\dot{X}_3(U)$, as follows. We compute the support associated to this new expression $\dot{X}_3(U)$, and all the projections with the Hilbert basis calculated for this case, 2.15, i.e., $\langle a_i^*, m_j \rangle$, where $i = 1, 2, 3, 4, 5, 6, 7, 8$, runs for the elements in the Hilbert basis and $j$ for the elements in the support.

$Supp(\dot{X}_3(U)) =$
$\{m_1 = (0, 0, 0, 0, 0, 1), m_2 = (0, 2, 2, 2, 0, 3), m_3 = (0, 0, 0, 2, 0, 1), m_4 = (0, 0, 1, 2, 0, 1)\}$.

We calculate all the projections associated to these faces, for the rays $m_1$, $m_2$, $m_3$ and $m_4$ ,with the elements of the Hilbert basis $a_j^*$, that is to say, all the scalar products, $\langle a_j^*, m_1\rangle$, $\langle a_j^*, m_2\rangle$, $\langle a_j^*, m_3\rangle$, $\langle a_j^*, m_4\rangle$.

$$\langle a_1^*, m_1\rangle = 0, \text{ and } \langle a_1^*, m_2\rangle = 0,$$
$$\langle a_2^*, m_1\rangle = 0, \text{ and } \langle a_2^*, m_2\rangle = 2$$
$$\langle a_3^*, m_1\rangle = 0, \text{ and } \langle a_3^*, m_2\rangle = 4$$
$$\langle a_4^*, m_1\rangle = 0, \text{ and } \langle a_4^*, m_2\rangle = 8.$$
$$\langle a_5^*, m_1\rangle = 0, \text{ and } \langle a_5^*, m_2\rangle = 0.$$
$$\langle a_6^*, m_1\rangle = -1, \text{ and } \langle a_6^*, m_2\rangle = 1.$$

For the vectors, $m_3$ and $m_4$, we get the minimal projections.

$$\langle a_1^*, m_3\rangle = 0, \text{ and } \langle a_1^*, m_4\rangle = 0,$$
$$\langle a_2^*, m_3\rangle = 0, \text{ and } \langle a_2^*, m_4\rangle = 0$$
$$\langle a_3^*, m_3\rangle = 0, \text{ and } \langle a_3^*, m_4\rangle = 1$$
$$\langle a_4^*, m_3\rangle = 2, \text{ and } \langle a_4^*, m_4\rangle = 4.$$
$$\langle a_5^*, m_3\rangle = 0, \text{ and } \langle a_5^*, m_4\rangle = 0.$$
$$\langle a_6^*, m_3\rangle = -1, \text{ and } \langle a_6^*, m_4\rangle = 0.$$

The minimal branch $\gamma$ which intersects the faces is defined by the projections, with $\langle a_6^*, m_3\rangle = -1$ and $\langle a_6^*, m_1\rangle = -1$, therefore the only terms to study the equation $\dot{X}_3(U)$ are in $m_1$ and $m_3$. For a deep study of bifurcation in dynamical systems and branches, see, [], [], [], [], thus, we have.

$$\dot{X}_3(U)_\gamma = c_1 * u_6 + c_3 * u_4^2 u_6. \tag{2.16}$$

We can notice in this example, the hilbert basis also help us to detect a dominant flux, detecting the minimal projections on faces of cone from S-system.

In the next sections, we only are interested in solving geometrical regions to bound kinetic orders.

### 3. Computing kinetic orders in Topological Environment.

To continuation we explain with details an algorithmic methodology for the bulding of local geometrical regions, and therefore to get bounds for kinetic orders in S-systems. Our main input from any genetic-metabolic network, will be the environment cone associated to an S-system to analyse.

**Algorithm:**

**Input:** *support*$(S)$**.** We take the exponent space or support from S-system to analyse, $(g_{1,p_1}, ..., g_{n,p_n})$ and $(h_{1,q_1}, ..., h_{n,q_n})$.

**Step 0: Constructing the environment cone associated to the support .** We locate the biomolecule $X_i$ associated to the S-system on the network, we consider the node neighbors, they are different kind of biomolecules, the kinetic orders are unknown. We construct stochastic matrices with arbitrary weights of kinetic orders, these weights are chosen in a random way among $0-9$ for any species, they are curated values involved in any biochemical process in metabolism, is to say this the biochemical environment around of $X_i$.

**Step 1: Constructing a closed cone** The vectors or rays of the cone defined in the input, they are open cones, as we shown below in the figure, we need closed cones by topological requirements, [], so that we can get closed cones, including the vector $(g_{1,p_1}, ..., g_{n,p_n}) - (h_{1,q_1,...,h_{n,q_n}})$ in the original cone, with it we continue to step 2.

**Step 2: Orthogonal enzyme kinetics $\sigma^\vee(S)$.** (To compute a big data-set of stochastic evironment dual cones.) We chosen the stochastic matrices of step 0. with the condition what the rows fullfil the orthogonality with the vectors in the step 1, i.e, $\langle (g_{1,p_1}, ..., g_{n,p_n}), (b_1, ..., b_n) \rangle = 0$, this is the simplest way to construct in a random way, arbitrary stochastic matrices. We give a python code to the generation of environment cones.

```
def Cone(n) :

r=0

while r ¡ n :

r=r+1

A=np.loadtxt(”Matrix_Cone_Sim1.txt”)

for j in range(4):

for i in range(3):

if A[j].item(i)!=0:

print(np.put(A[j],[i],[0]))

elif A[j].item(i)==0 :

print(np.put(A[j],[i],[np.random.randint(1,9)]))

print(A)

np.savetxt( 'MATRIX_DUAL_CONE_X1.txt'.format(r), A, delimiter = ”,”, newline ='
', header =' intmatH[4][3] =' .format(r), fmt = ”
else:

print(np.put(A[j],[i],[-1]))

r=r+1
```

**Step 3: Hilbert basis** $H_{i\sigma^\vee}$ (To compute of Hilbert basis in Singular computer algebra programm for each stochastic cones ). We compute the Hilbert basis of matrices constructed in setp 2. with following script in Singular, [].

$inti;$
$intj;$
$intt;$
$for(i = 1; i <= 4; i = i + 1)$
$conec(i) = coneViaPoints(L[i]);$
cone d(i)=dualCone(c(i));

matrix H(i)=hilbertBasis(d(i));

**Step 5: Unimodular Hilbert matrices** $[\cap H_i]_{mxm}$ We only get uni-modular matrices (with determinant $\pm 1$) from the intersection $_i$ of all the set of Hilbert basis computed in the step 3., implying to get a local geometry for kinetic orders, we do that with the followig script.

$for(j = 1; j <= 4; j = j + 1)$

H(j);

for(t=1;t¡=nrows(H(j))-2;t=t+1)

intvec k1=nrows(H(j))-t-1;

intvec k2=nrows(H(j))-t;

intvec k3=nrows(H(j))-t+1;

intvec K=k1,k2,k3;

matrix B(t)=submat(H(j),K,1..3);

if (det(B(t))==1 —— det(B(t))==-1)

matrix SH(t)=B(t);

print(SH(t));

else

return t;

**Step 6: Using the condition of fix points from unimodular Hilbert matrix**. With the elements of the unimodular Hilbert basis, we compute the constrains of inner products $\langle (g_{1,p_1}, ..., g_{n,p_n}), (h_1, ..., h_n) \rangle >= 0$, this process is done for each vector of step 1, $(g_{1,p_1}, ..., g_{n,p_n})$, and for each vector in the Hilbert basis, $(h_1, ..., h_n)$ .

**Step 7: Set of inequalities built from the condition of fix points.** The constraints computed in the sptep 6, are a system of inequalities must be valids for that the limits in the formula, 2.7, can exist, the kinetic orders are icognites of system, when we plot these inequalities in the plane, we get regions that bounds numerical values for kinetic orders.

**Output: Geometrical regions from step 7 bounds kinetic orders in S-sytems**.

### 4. Applications in a toy genetic-metabolic network.

**Description of the gene circuit.**

To continuation we will expose the main example of our work. We analyse the following gene-metabolic circuit, fig. 5, it consists of a negative loop of regulation of two regulator genes, the first gene activates the expression of the second, and this one repress the expression of the first gene. We start the analysis considering that the kinetic parameters, $\alpha_{max}$, $\alpha_{min}$, $\alpha_2$, $\beta_2$, $\beta_1$ and kinetic orders, $p_1$, $p_2$, $p_3$, and $p_4$ are unknown. We will determine their values using Design space toolbox for the first parameters, and for the seconds parameters we utilize the algorithm described here.

TOY_CIRCUIT.png

FIGURE 5. System of two regulators.

We give the expressions for the rate laws of the two genes, it is shown as we can see below,

$$\dot{X}_1 = \alpha_{min} D^{-1} + \alpha_{max} \left( \frac{X_2}{K_A} \right)^{p_1} D^{-1} - \beta_1 X_1^{p_2}, \tag{4.1}$$

$$\dot{X}_2 = \alpha_2 X_1^{p_2} - \beta_2 X_2^{p_4}, \tag{4.2}$$

$$f = 0 = 1 + K_A^{-p_1} X_2^{p_1} - D. \tag{4.3}$$

**4.1. Exponent space and Support of S-systems associated.** The supports obtained for each expression above in lexicographic order are given by:

$$\text{supp}(\dot{\text{X}}_1) = \begin{pmatrix} p_2 & 0 & 0 \\ 0 & p_1 & 0 \\ 0 & 0 & -1 \end{pmatrix} \ (4.4)$$

$$\text{supp}(\dot{\text{X}}_2) = \begin{pmatrix} p_3 & 0 & 0 \\ 0 & p_4 & 0 \end{pmatrix} \ (4.5)$$

$$\text{supp(f)}= \begin{pmatrix} 0 & p_1 & 0 \\ 0 & 0 & 1 \end{pmatrix} \ (4.6)$$

**Cones associated**. We take the substration row by row of these matrices to contruct closed cones as points the step 1, we have.

$$\text{Con}(\dot{\text{X}}_1) = \begin{pmatrix} p_2 & 0 & 0 \\ p_2 & -p_1 & 1 \\ 0 & p_1 & 0 \\ 0 & 0 & -1 \end{pmatrix} \ (4.7)$$

$$\text{Con}(\dot{\text{X}}_2) = \begin{pmatrix} p_3 & 0 & 0 \\ p_3 & -p_4 & 0 \\ 0 & p_4 & 0 \end{pmatrix} \ (4.8)$$

$$\text{Con(f)}= \begin{pmatrix} 0 & p_1 & 0 \\ 0 & p_1 & -1 \\ 0 & 0 & 1 \end{pmatrix} \ (4.9)$$

#### 4.2. Computing stochastic environment cones. Matrices computed from files.

We did simulations with the code in python and sigular described by the algorithm given in the fisrt part of the section,we computed 40 thousands miles stochastic evorinoment matrices (environment cones), 20 thousand miles for each cone $Con(\dot{X}_1)$ and $Con(f)$, respectevely. The cone $Con(\dot{X}_2)$ always has associated the identity matrix as Hilbert basis, in thar case is not required simulations, we simply intersect this matrix with the other ones.

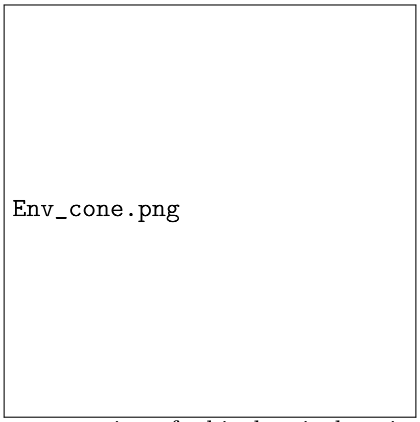


Figure 6. Representation of a biochemical environment cone for a biomolecule associated to an S-system.

These are some example generated by the algortihm, in the part of supplementary material we add the files with the set of matrices simulated.

$intmatH839[4][3] = 5, 0, 2, 4, 7, 0, 0, 5, 0, 0, 0, 6;$

intmat H105[4][3]=1,0,6,7,5,0,0,3,0,0,0,4;

intmat H1755[4][3]=5,0,6,4,2,0,0,7,0,0,0,2;

intmat H1877[4][3]=4,0,3,3,3,0,0,7,0,0,0,2;

#### 4.3. Computing Hilbert basis in evironment cones. Set of unimodular Hilbert basis.

**Computing unimodular Hilbert Basis.** We show one of the Hilbert basis simulated by the algorithm, here the, 1, 3 and 5 rows marked with starts in the matrix, are invariant vectors that appears with high score of probabilty (analysis with p-value) through of all set of simulations.

EXAMPLE:

h(7)[1,1]=1******

h(7)[1,2]=0******

h(7)[1,3]=0******

h(7)[2,1]=-6

h(7)[2,2]=6

h(7)[2,3]=5

h(7)[3,1]=0******

h(7)[3,2]=0******

h(7)[3,3]=1******

h(7)[4,1]=0

h(7)[4,2]=1

h(7)[4,3]=0

h(7)[5,1]=-1*******

h(7)[5,2]=1

h(7)[5,3]=1 *******

**4.4. Geometrical regions for kinetic orders.** From the invariant unimodular matrix we can map these vectors into of invariant geometrical regions, those regions bounds values for kinetic orders for the equations, 4.1 and 4.2,

$$\text{unimodular(H)=} \begin{pmatrix} 1 & 0 & 0 \\ 0 & 1 & 0 \\ -1 & 1 & 1 \end{pmatrix} \ (4.10)$$

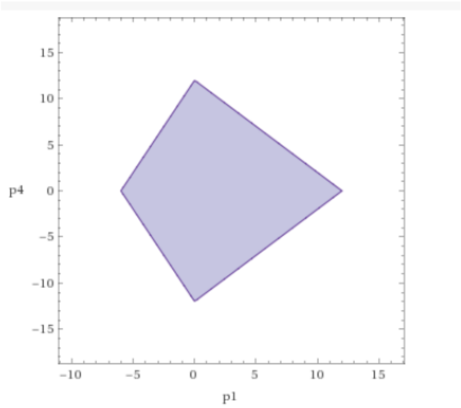


FIGURE 7. Plot for the region that bounds the kinetic orders, $p_1$ and $p_4$.

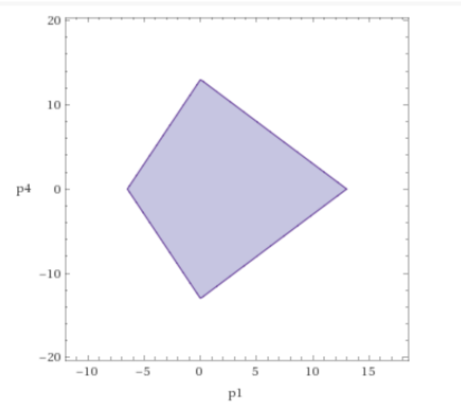


FIGURE 8. Plot for the region that bounds the kinetic orders, $p_2$ and $p_3$.

The figures 7 and 8. are plots showing closed regions that contains ordered pairs of coordinates, $(p_1, p_4)$ and $(p_2, p_3)$, that regions are mapped from unimodular Hilbert matrix 4.10, any pairs into the regions, are valid values to solve the differential equations, 4.1 and 4.2 in our example, to continuation we take some values to solve dyanmics with Design space toolbox.

**4.5. Dynamics and quantitative phenotypes with Desig-Space Toolbox v2. Analysis Miguel and Mike**.

### 5. Discussion.

We have shown new topological properties in biochemical networks, exploting the characteristics of Power Law and S-systems modeling, we notice mainly the monomial map property, $g_{ij} \longrightarrow X_j^{gij}$. we take advantage of this map analysing

only the exponent space in S-systems, as consequence we focus our analysis to kinetic orders by means of computational algebraic geometry techniques to find geometrical basis to bound the values of these parameters. This technique let us to solve big set of differential equations. The new issues to solve with this technique, are : when are valid these regions to give values to kinetic oredrs?, what geometrical regions are mathematical artifacts and which are regions with biological information for the netwok analysis.
Another of the advantages of the methodology presented here, is the possibilty of modelling for dynamics in large-scale networks in metabolism. The property of monomial map has the formal justyfication to module netowrks through the mathemtical concept of **toric variety**, they are abstract mathematical objects, but they are capable to give information in modules, to analyse module by module thorugh varieties and after to glue thogether in a big toric variety. Pherphaps, the information that provides one variety is not very informative for the biochemical network, but the whole pieces or varierties glueded togheter let us see the complete dynamics in large-scale networks.

### 6. Conclusions.

We do emphasis in the information given by the relationship given by, $g_{ij} \longrightarrow X_j^{gij}$, as an alternartive model to gain knowledge in the complex molecular connection, genotype-phenotype. With this map we have obtained new topological properties for analysis with S-systems through the fix points on torus and Hilbert basis, the consequence for genetic-metabolic networks, are: to find out an particular steady-state for genotypes, an equivalence to the classical steady-state for phenotypes, with this new knowledge we can compute geometrical regions to bound values for unknown kinetic orders, for any network, using the modular concept of algebraic **variety**, as an artifice to solve big networks, but with a general biological relevance in the dynamics solution for genetic-metabolic networks.

### 7. Acknowledgment

We acknowledge funding from UNAM and from the National Institutes of Health (grant number 5R01GM110597-03).

Department of Molecular Biology and Biotechnology, Institute of Biomedical Research UNAM, Circuito Universitario 04510 CDMX, Mexico
