## Supplementary material for "Towards Modeling in Large-Scale Genetic and Metabolic Networks": Latex_Manuscript: Topological_Env_kineticorders.tex

%% filename: amsproc-template.tex
%% version: 1.1
%% date: 2014/07/24
%%
%% American Mathematical Society
%% Technical Support
%% Publications Technical Group
%% 201 Charles Street
%% Providence, RI 02904
%% USA
%%      (800) 321-4267 (USA and Canada only)
%%
%% 
%% Copyright 2008-2010, 2014 American Mathematical Society.
%% 
%% This work may be distributed and/or modified under the
%% conditions of the LaTeX Project Public License, either version 1.3c
%% of this license or (at your option) any later version.
%% The latest version of this license is in
%%   http://www.latex-project.org/lppl.txt
%% and version 1.3c or later is part of all distributions of LaTeX
%% version 2005/12/01 or later.
%% 
%% This work has the LPPL maintenance status `maintained'.
%% 
%% The Current Maintainer of this work is the American Mathematical
%% Society.
%%
%% ====================================================================

%     AMS-LaTeX v.2 template for use with amsproc
%
%     Remove any commented or uncommented macros you do not use.

\documentclass{amsproc}

\newtheorem{theorem}{Theorem}[section]
\newtheorem{lemma}[theorem]{Lemma}

\theoremstyle{definition}
\newtheorem{definition}[theorem]{Definition}
\newtheorem{example}[theorem]{Example}
\newtheorem{xca}[theorem]{Exercise}

\theoremstyle{remark}
\newtheorem{remark}[theorem]{Remark}

\numberwithin{equation}{section}
\usepackage{float}
\usepackage{graphicx}
\usepackage{verbatim}
\usepackage{amsmath}
\usepackage{xcolor}

\begin{document}

\title[]{Towards Modeling in Large-Scale Genetic and Metabolic Networks.}
%\title{Topological Environment of genetic-metabolic Networks:
%Towards a modelling for Large-Scale Networks.}
\author{Marco Polo Castillo-Villalba}
\address{Department of Molecular Biology and Biotechnology, Institute of Biomedical Research UNAM, Circuito Universitario 04510 CDMX, Mexico}
%\curraddr{}
%%%%***\author{Miguel \'{A}. Valderrama-G\'{o}mez}
%%%%***\address{Department of Microbiology and Molecular Genetics, University of California, Davis, CA 95616, USA}
%%%%***\author{Michael A. Savageau}
%%%%%***\address{Department of Biomedical Engineering, University of California, Davis, CA 95616, USA}
%%%***\author{Julio Collado-Vides}
%%%***\address{Program of Computational Genomics, Center for Genomic Sciences UNAM, C.P. 62210, Cuernavaca, Morelos, Mexico.}
%%**\author{Luis Mendoza} 
%%%***\address{Department of Molecular Biology and Biotechnology, Institute of Biomedical Research UNAM, Circuito Universitario 04510 CDMX, Mexico}

\subjclass[2010]{Primary }

\keywords{Topological Environment, S-systems, toric geometry, quantitative phenotype, kinetic orders, tryptophan gene-metabolic network.}

\date{7/8/2017}


%\begin{abstract}

%The mathematical modeling for large-scale genetic and metabolic networks is even an open topic for research with deep implications in the signal biochemical systems such as organisms prokaryotes as eukaryotes. We have used the biochemical modeling (Power Law formalism) widely developed by M.A. Savageau in small networks to elucidate principle of design found in the bacterial cells. this modeling describes events such as gene regulation as metabolic fluxes, both biological processes are well characterized mainly by two sets of parameters, kinetic parameters, and kinetic orders. The limitation to describe larger networks is the lack of that parameters. Once provided any family of those parameters is possible to predict the second one. We propose a mathematical technique to predict values to the family associated with the space of kinetic orders. They are the exponents in the power law representation and the kinetic parameters are the coefficients of the model. We have proved that this exponent space of kinetic orders is well-known in toric geometry and it can be modeled by algebraic varieties. To model a large network only we associate a variety in each polynomial and then we can glue together the varieties by geometrical techniques.  
%\end{abstract}

%\maketitle
%\section{Introduction}


  %%%%%%%%%%%%%%%%%%%%%%%%%%%%%%%%%%%%%%%%%%%%%%%%%%%%%%%%%%%%%%%%%%%%%%%%%%%%%
%%%%%%%%%%%%%%%%%%%%%%%%nueva secccion%%%%%%%%%%%%%%%%%%%%%%%%%%%%%%%%%%%%%%%%%%%
%\section{Theoretical background of Biochemical networks as Toric S-systems.}

%One of the main ideas behind analysis of biochemical system is the balance of mass in the reactions, in additon Micheles-Menten rate law, is an of the concept fundamentals in biochemical systems, but in the last decades this law has been improved to give more biochemical details of the biomolecule participatiyng in the network, for example now we can consider any arbitrary order of reaction, as to consider the number of binding sites of operator region in genes, or an activation parameter of Hill's function. It is the case of the theory of power law formalism, \cite{}, where considering this parametric extension, in this modelling is possible to include matabolism and gene regulation togheter. The formal expression of this model is presented below.\\
  
%\begin{equation}
 %\dot{X}_{i}= \sum_{k=1}^{p_{i}} \alpha_{ik}prod_{j=1}^{m}X_{j}^{g_{ijk}} - \sum_{k=1}^{q_{i}} \beta_{ik}\prod_{j=1}^{m} X_{j}^{h_{ijk}},\: i=1,...,n.
%\end{equation}

%Here, the kinetic parameters, $ \alpha_{ik} $, $ \beta_{ik}  $, capture information of constant rate laws of production and degradation repectevely, of any species or biomolecule in the network, the kinetic orders $g_{ijk}  $, $h_{ijk}  $, capture genotypic information, from bindings sites and kinetic orders of catalysed reactions, as we have commented at first. Hence, the expression 2.1, has captured all information to analyse genetic-metabolic networks.   

%\begin{definition}\label{Definition 3.2} (\textbf{S-systems}). We define a number of combinations of dominant terms in 2.1, given by: \[ \prod_{j=1}^{m} p_{j}q_{j} ; \]  partitions of the space of concentrations $(X_{1},...,X_{n}) $, to each partition we call it, a dominant term or \textbf{S-system}, \cite{MASavageauVoit1987}, \cite{MASavageau1976}, \cite{LomnitzSavageau2013}, \cite{LomnitzSavageau2014}, \cite{LomnitzSavageau2015}; as shown below,
%\begin{equation}
%\dot{X}_{i}=  \alpha_{ip_{i}} \prod_{j=1}^{m} X_{j}^{g_{ip_{i}}} -  \beta_{iq_{i}} \prod_{j=1}^{m}X_{j}^{h_{iq_{i}}}, \, i=1,...,n.\\
%\[ 0=  \alpha_{ip_{t+1}} \prod_{j=1}^{m}X_{j}^{g_{ip_{t+1}}} -  \beta_{iq_{t+1}} \prod_{j=1}^{m}X_{j}^{h_{iq_{t+1}}}, \, t=0,...,m,
%\end{equation}
%\end{definition}\

%In the context of this formalism, an S-system defines a quantitative phenotype and also corresponds to a reality of molecular biology, not only as mathematical artifacts, clearly they are rate of molecular concentrations computed by this gradient.
\begin{abstract}
Mathematical modeling of large-scale genetic and metabolic networks remains an open area of research with profound implications for understanding biochemical signaling systems in both prokaryotic and eukaryotic organisms. In this work, we extend the biochemical modeling framework based on the \textbf{Power Law formalism}, originally developed by M.A. Savageau to elucidate design principles in small bacterial networks. This formalism captures gene regulation and metabolic fluxes through two fundamental sets of parameters: \textit{kinetic parameters} and \textit{kinetic orders}. The kinetic parameters determine the magnitudes of fluxes, while the kinetic orders appearing as exponents in the power-law representation encode the regulatory structure of the system. We propose a mathematical technique to characterize the entire family of systems associated with the space of kinetic orders. We show that this exponent space possesses a well-defined structure in \textbf{toric geometry} and can therefore be modeled using \textbf{algebraic varieties}. To represent large-scale networks, we associate a toric variety with each polynomial equation of the system and then use geometric techniques to coherently combine these varieties into a global structure, or \textbf{fan}. This construction enables a modular and geometrically consistent representation of complex biochemical networks.

%Mathematical modeling of large-scale genetic and metabolic networks remains an open area of research with deep implications for signal biochemical systems in both prokaryotic and eukaryotic organisms. We have applied the biochemical modeling approach (Power Law formalism) widely developed by M.A. Savageau for small networks to elucidate the design principles found in bacterial cells. This modeling framework describes events such as gene regulation and metabolic fluxes, both of which are characterized mainly by two sets of parameters: kinetic parameters and kinetic orders, the kinetic parameters are analysed from power-law formalism. In this work, is propose a mathematical technique to characterize the family associated with the space of kinetic orders. These orders represent the exponents in the power law representation, while the kinetic parameters correspond to the coefficients of the model. We demonstrate that this space of kinetic orders is well-known in toric geometry and can be modeled using algebraic varieties. To model a large network, we associate a variety with each polynomial and then use geometric techniques to combine these varieties coherently. With the aim to model a whole network gluying varieties together in a fan.

\end{abstract}

\maketitle
\section{Introduction}

%%%%%%%%%%%%%%%%%%%%%%%%%%%%%%%%%%%%%%%%%%%%%%%%%%%%%%%%%%%%%%%%%%%%%%%%%%%%%
%%%%%%%%%%%%%%%%%%%%%%%%% New Section %%%%%%%%%%%%%%%%%%%%%%%%%%%%%%%%%%%%%%%
\section{Theoretical Background: Biochemical Networks as Toric S-systems}

One of the main ideas behind the analysis of biochemical systems is the conservation of mass in reactions. In addition, the Michaelis–Menten rate law is one of the fundamental concepts in biochemical systems. However, in recent decades, this law has been refined to provide more detailed biochemical information about the biomolecules participating in the network. For example, it is now possible to consider reactions with arbitrary orders, such as the number of binding sites in gene operator regions or activation parameters from Hill functions. 

This is the basis of the Power Law formalism \cite{MASavageau1976}, which, through parametric extensions, allows for the simultaneous modeling of both metabolism and gene regulation. The formal expression of this model is given below:

\begin{equation}
 \dot{X}_{i}= \sum_{k=1}^{p_{i}} \alpha_{ik} \prod_{j=1}^{m}X_{j}^{g_{ijk}} - \sum_{k=1}^{q_{i}} \beta_{ik}\prod_{j=1}^{m} X_{j}^{h_{ijk}}, \quad i=1,\ldots,n.
\end{equation}

Here, the kinetic parameters $\alpha_{ik}$ and $\beta_{ik}$ capture the rate constants for production and degradation, respectively, of each species or biomolecule in the network. The kinetic orders $g_{ijk}$ and $h_{ijk}$ represent genotypic information, such as binding sites and the kinetic orders of catalyzed reactions or multimeric regulation sub units in protein structures. Therefore, Equation (2.1) captures all the information required to analyze genetic–metabolic networks.

\begin{definition}\label{Definition 3.2} 
(\textbf{S-systems}). We define the number of combinations of dominant terms in Equation (2.1) as:
\[ \prod_{j=1}^{m} p_{j}q_{j}, \]
which partitions the space of concentrations $(X_{1}, \ldots, X_{n})$. Each partition corresponds to a dominant term, or \textbf{S-system}, as defined in \cite{MASavageauVoit1987}, \cite{MASavageau1976}, \cite{LomnitzSavageau2013}, \cite{LomnitzSavageau2014}, \cite{LomnitzSavageau2015}. This is expressed as follows:
\begin{equation}
\dot{X}_{i}=  \alpha_{ip_{i}} \prod_{j=1}^{m} X_{j}^{g_{ip_{i}}} -  \beta_{iq_{i}} \prod_{j=1}^{m}X_{j}^{h_{iq_{i}}}, \quad i=1,\ldots,n.
\end{equation}
\end{definition}

In the context of this formalism, an S-system defines a quantitative phenotype and corresponds to a biological reality in molecular biology. These systems are not merely mathematical artifacts; they represent rates of change in molecular concentrations derived from this formal gradient.

\subsection{Toric S-systems}

In this section, we highlight the importance of studying the exponents $g_{ip_{i}}$ and $h_{iq_{i}}$ in an S-system. We assert that the monomial map $g_{ip_{i}} \longmapsto X_{j}^{g_{ip_{i}}}$ represents a \textbf{Genotype–Phenotype} relationship with properties of great interest for further study. This central idea forms the foundation of our research. Below, we briefly summarize the definitions introduced in well-known works such as \cite{LomnitzSavageau2013}, \cite{MASavageauVoit1987}, \cite{MASavageau1976}, and \cite{LomnitzSavageau2013}.\\

\begin{definition}\label{Definition 3.2.1} 
\textbf{(Exponent Space or Support)}  
For an S-system defined by $\dot{X}_{i}$, we define the \textit{exponent space} of $\dot{X}_{i}$, also formally known as the \textit{support} of a polynomial, as:
\begin{equation}
\mathrm{supp}(\dot{X}_{i}) = \left\lbrace (g_{i,p_{1}}, \ldots, g_{i,p_{n}}), (h_{i,q_{1}}, \ldots, h_{i,q_{n}}) \right\rbrace.
\end{equation}

Using the support of any S-system $\mathrm{supp}(\dot{X}_{i})$, we can construct all linear combinations from the two vectors in the support, $(g_{1,p_{1}}, \ldots, g_{n,p_{n}})$ and $(h_{1,q_{1}}, \ldots, h_{n,q_{n}})$. The set of all such vector combinations defines the \textbf{cone} associated with $\dot{X}_{i}$, as described in \cite{GEwald1993}.
\end{definition}

\begin{definition}\label{Definition 3.2.2} 
\textbf{(Cone)}  
Let $\dot{X}_{i}$ be an S-system as defined above. We define the cone associated with it as:
\begin{equation}
\sigma = \mathrm{Con}\left( \mathrm{supp}(\dot{X}_{i}) \right).
\end{equation}
This cone represents the linear space generated by all combinations of the vectors in the support space, including addition, subtraction, and scalar multiplication. If the vectors consist only of integer entries, the cone is called a \textit{lattice cone} \cite{GEwald1993}.\\
\end{definition}

\begin{definition}\label{Definition 3.2.3} 
\textbf{(Dual Cone)}  
Let $\sigma \subseteq \mathbb{Z}^{n}$ be a lattice cone. The dual cone $\sigma^{\vee}$ is defined as:
\begin{equation}
\sigma^{\vee} = \left\lbrace v \in \mathbb{Z}^{n} \; : \; \langle v, \sigma \rangle \geq 0 \right\rbrace,
\end{equation}
as described in \cite{GEwald1993}.
\end{definition}

\begin{definition}\label{Definition 3.2.4} 
\textbf{(Monoid)}  
By intersecting the dual cone with the $n$-dimensional integer lattice, we obtain:
\[
\sigma^{\vee} \cap \mathbb{Z}^{n},
\]
which we call a \textbf{monoid}. This monoid, denoted by $M$, has rank $n$ and is isomorphic to $\mathbb{Z}^{n}$ \cite{GEwald1993}.\\
\end{definition}

\textbf{Lemma (Gordan’s Lemma and the Hilbert Basis).}  
Given a cone $\sigma$, the intersection of its dual with the $n$-dimensional integer space,
\[
\sigma^{\vee} \cap \mathbb{Z}^{n},
\]
forms a monoid that has a finite set of generators. This generating set is called the \textbf{Hilbert basis} \cite{BSturmfels1996}, \cite{GEwald1993}.\\

\begin{definition}\label{Definition 3.2.5} 
\textbf{(Fan or topological environment)}  
Given a collection of dual cones 
\[
\left\lbrace \sigma^{\vee}_{1}, \ldots, \sigma^{\vee}_{n} \right\rbrace \subseteq \mathbb{Z}^{n},
\]
we define the \textit{fan} $P = \bigcup_{i=1}^{n} \sigma^{\vee}_{i}$ as the union of all these cones. We denote this fan by $\Sigma \subseteq \mathbb{Z}^{n}$ \cite{GEwald1993}.\\
\end{definition}

Finally, we construct a coordinate polynomial ring for S-systems.  
Let $f = \dot{X}_{i}$ be an S-system defined for any $i$-th species in a metabolic network. Given the support of this polynomial $\mathrm{supp}(f)$ and its associated cone $\sigma$, we define the polynomial ring:
\[
R_{\sigma} = \left\lbrace f \; \middle| \; \mathrm{supp}(f) \subseteq \sigma \right\rbrace.
\]
In other words, the exponents of an S-system (i.e., the kinetic orders) are embedded in a geometric space. We will explore new topological properties that arise from this geometric object in the following sections.\\


%\subsection{Toric S-systems.}

%In this section, we remark on the importance to study of the exponents $g_{ip_{i}}  $ and $h_{iq_{i}}  $  in an S-system, and we do the assertion that the monomial map $g_{ip_{i}} \longmapsto X_{j}^{g_{ip_{i}}}  $ is a relationship \textbf{Genotype-Phenotye} with interesting properties to study, and this central idea, is the fundamental ground of our research. We summarize in brief definitions given in well-known works, \cite{LomnitzSavageau2013}, \cite{MASavageauVoit1987}, \cite{MASavageau1976}, \cite{LomnitzSavageau2013}.\\

%\begin{definition}\label{Definition 3.2.1} \textbf{(Exponent-space or support)} For an S-system given $ \dot{X}_{i} $, we define the exponent-space of $\dot{X}_{i}  $ or formally called support of a polynomial, as,\\
%\begin{equation}
%supp(\dot{X}_{i} )= \left\lbrace (g_{i,p_{1}},...,g_{i,p_{n}}), (h_{i,q_{1}},...,h_{i,q_{n}} ) \right\rbrace 
%\end{equation}

%With the support of the any S-system $ supp(\dot{X}_{i} ) $, we can build all the linear combinations from the two vectors in the support, $ (g_{1,p_{1}},...,g_{n,p_{n}}) $ and $ (h_{1,q_{1},...,h_{n,q_{n}}} ) $, this set of combinations of vectors define the \textbf{Cone} associated to $ \dot{X}_{i} $, see \cite{GEwald1993}.

%\begin{definition}\label{Definition 3.2.2} \textbf{(Cone)}. Let  $\dot{X}_{i} $ be an S-system defined as above, we define the cone associated with it, as follows,\\

%\begin{equation}
%\sigma =Con\left( supp(\dot{X}_{i} ) \right) .
%\end{equation}


%It is the linear space defined by all combinations for the vectors of support space, including addition, subtraction and scalar multiplication of vectors, if the vectors consist only of integer entries, is called a lattice-cone, see \cite{GEwald1993}.\\

%\begin{definition} \label{Definition 3.2.2} \textbf{(Dual cone)}.
%\begin{equation}
%Let  $\sigma \in \mathbb{Z}^{n}$ be a lattice-cone \sigma^{\vee} = \lbrace \forall v \in \mathbb{Z}^{n} :  \:  < v , \sigma >  \quad > = \quad 0 \rbrace, see \cite{GEwald1993}.
%\end{equation} 

%\begin{definition}\label{Definition 3.2.2} \textbf{(Monoid)}.
%ow we intersect the cone with the $ n- $dimensional integer space, we write, $\sigma^{\vee} \cap \mathbb{Z}^{n} $, we call to this set as a \textbf{Monoid} or as a set $M$ of rank $n$ or isomorphic to $ \mathbb{Z}^{n} $, see \cite{GEwald1993}. \\

%\textbf{Lemma} \textbf{(Gordan's Lemma)}.\textbf{Hilbert basis} .  Given a cone $ \sigma $ and we take the intersection with the $ n- $dimensional integer space $\sigma^{\vee} \cap \mathbb{Z}^{n} $, then this monoid, has a finite set of generators, this set of generators is called, the \textbf{Hilbert basis}, see \cite{BSturmfels1996}, \cite{GEwald1993}.\\
%\begin{definition}\label{Definition 3.2.2} \textbf{(Fan)}. Given a set of dual-cones
%$\left\lbrace \sigma^{\vee}_{1},...,\sigma^{\vee}_{n} \right\rbrace \:\subseteq \:\mathbb{Z}^{n} $. We define the fan $ P =\sum=\bigcup^{n}_{i=1}\sigma^{\vee}_{i} $, which is the union of all these cones. We denote to this fan as $\sum \:\subseteq \:\mathbb{Z}^{n} $, \cite{GEwald1993}\\

%We build a coordinate polynomial ring of S-systems. Let be $ f = \dot{X}_{i} $ an S-system defined for any $ i- $esim specie in the metabolic network, given the support of this polynomial $ supp(f) $ and the cone associated with this set, $ \sigma $, we define a polynomial ring, as follows, $ R_{\sigma}= \left\lbrace f \: \mid \: supp(f) \:\subseteq \: \sigma  \right\rbrace   $. In other words, the exponents of an S-system or kinetic orders are embedded in a geometrical space, we will exploit new topological properties on this new geometrical object, as we can see.\\

%%%%%%%%%%%%%%%%%%TORIC IDEAL %%%%%%%%%%%%%%%%%%%%%%%%%%%%%%
\begin{definition}\label{Definition 3.2.2} \textbf{(Toric Ideal)}. A toric ideal is the set of prime binomials defined by $ I = <X^{A} - X^{B} | H*(A - B)=0 >$, where $H$ is the \textit{Hilbert basis} of the polynomials generated from $I$, \cite{BSturmfels1996}, \cite{DCoxLittleSchenck2010}. 
%%%%%%%%%%%%%%%%%%%%%%%%%%%%%%%%%%%%%%%%%%%%

\begin{definition}\label{Definition 3.2.2} 
\textbf{(Toric Variety)}.  
We define a \textbf{toric variety} associated with a polynomial $f$ as follows:
\begin{equation}
X_{\sigma} = \mathrm{Spec}\left( R_{\sigma} \right) = \left\lbrace f_{1}, \ldots, f_{k} \in R_{\sigma} \right\rbrace.
\end{equation}

Here, the notation $\mathrm{Spec}\left( R_{\sigma} \right)$ denotes the set of all prime polynomials (i.e., non-divisible polynomials) $\{ f_{1}, \ldots, f_{k} \}$ in the ring $R_{\sigma}$ for some integer $k$, see \cite{GEwald1993}. 
\end{definition}

\begin{definition}\label{Definition 3.2.3} 
\textbf{(Environment Cone)}.  
Let $f$ be an S-system representing the rate law for the change in concentration of any species in a gene–metabolic network.  
We compute the support of $f$ and the cone associated with it, $\sigma \subseteq \mathbb{Z}^{n}$.  
Since $ \mathrm{supp}(f) \subseteq \sigma $, we define the \textbf{environment cone} associated with $f$ as the dual cone $\sigma^{\vee}$ of $\sigma$.  
Thus, we have the inclusion relation:
\[
\mathrm{supp}(f) \subseteq \sigma \subseteq \sigma^{\vee}.
\]
\end{definition}

\textbf{Relevance in Enzyme Kinetics.}  
Up to this point, we have equipped the set of vectors associated with the kinetic orders of an S-system with new geometric properties.  
We have defined cones, polynomial rings, and toric varieties.  
The most important result arises from \textbf{Gordan’s Lemma} \cite{GEwald1993}, which guaranties the existence of a generator basis (the \textbf{Hilbert basis}) for the entire kinetic order space defined for any species in the network.  
Next, we will explore how to exploit the properties induced by this Hilbert basis.\\

% -------------------------------------------------------------------------
% (Optional future section commented)
% -------------------------------------------------------------------------
% \textbf{Fixed Points on the Torus.}  
% There exist several algorithms for computing the Hilbert basis \cite{}, and once the set of generators 
% $H = \{h_{1}, \ldots, h_{k}\}$ for a cone $\sigma = \mathrm{Con}(b_{1}, \ldots, b_{r})$ is obtained, 
% where each vector $b_{j}$ defines a face, we can identify the fixed points on the torus or toric variety 
% through the following formula \cite{}:
% \begin{equation}
% X_{b_{j}} = \lim_{Z \to 0} Z^{\langle b_{j}, h_{i} \rangle} = (\lim_{Z \to 0}Z^{\langle b_{j}, h_{1} \rangle}, \ldots, \lim_{Z \to 0}Z^{\langle b_{j}, h_{k} \rangle}).
% \end{equation}
% These limits exist if and only if the following constraints hold:
% \[
% \langle b_{j}, h_{1} \rangle \geq 0, \; \langle b_{j}, h_{2} \rangle \geq 0, \ldots, \langle b_{j}, h_{k} \rangle \geq 0,
% \]
% for all relevant integers $j$ and $k$. 
% In this formulation, the vectors $b_{j}$ correspond to the kinetic orders associated with the support of a given S-system $f$. 
% This set of constraints, together with those for dominant fluxes, determines admissible values for kinetic orders within the algorithm described in this work.\\
%
% \textbf{Assertion.}  
% Classical fixed points correspond to steady states defined at the phenotype level. 
% In contrast, the fixed points on the torus encode information derived from the genotype, 
% which emerges naturally from the monomial map $g_{ip_{i}} \longmapsto X_{j}^{g_{ip_{i}}}$ studied above.\\

\textbf{Toric S-systems and Dominant Fluxes.}\\

Given the Hilbert basis $H = \{h_{1}, \ldots, h_{k}\}$, we represent it in matrix form considering each $h_{i}$ as a row vector:
\[
h_{1} = (u_{11}, u_{12}, \ldots, u_{1n}), \quad 
h_{2} = (u_{21}, u_{22}, \ldots, u_{2n}), \quad \ldots, \quad
h_{k} = (u_{k1}, u_{k2}, \ldots, u_{kn}).
\]

\begin{center}
$H  =   
\begin{pmatrix}
u_{11}&u_{12}&\cdots&u_{1n}\\
u_{21}&u_{22}&\cdots&u_{2n}\\
\vdots&\vdots&\ddots&\vdots\\
u_{k1}&u_{k2}&\cdots&u_{kn}
\end{pmatrix}$
\end{center}

We can use this matrix to represent the variables $X_{i}$ in a new coordinate system using the following monomial transformation:
\[
u_{ij} \longmapsto w_{i}^{u_{ij}},
\]
as discussed in \cite{GEwald1993}, \cite{BSturmfels1996}, and \cite{DCoxLittleSchenck2010}.  
This transformation reveals dynamical properties encoded within the S-systems through the Hilbert basis \cite{BSturmfels1996}, \cite{Henk}.  

\begin{equation}  
\begin{matrix}
X_{1} &=& w_{1}^{u_{11}} w_{2}^{u_{12}} \cdots w_{n}^{u_{1n}},\\
X_{2} &=& w_{1}^{u_{21}} w_{2}^{u_{22}} \cdots w_{n}^{u_{2n}},\\
\vdots & & \vdots\\
X_{k} &=& w_{1}^{u_{k1}} w_{2}^{u_{k2}} \cdots w_{n}^{u_{kn}}.
\end{matrix}
\end{equation}

This matrix representation allows embedding all S-system equations into a new system of monomial coordinates.  
Hence, we extend the analysis of molecular networks toward large-scale genetic and metabolic systems.\\

\begin{theorem}\label{Theorem} 
\textbf{(Toric S-system)}.  
If we substitute the above change of coordinates into Equation (2.2), the original S-system equations, we obtain the following expression, valid for all $i = 1, \ldots, n$:\\

\textbf{Proof.}  
Given an S-system representation of molecular networks as in Equation (2.2), and applying Hironaka’s theorem together with the algorithmic construction of the Hilbert basis (a finite number of times) to the S-system equations \cite{MFAtiyah1970}, \cite{TFukui2000}, \cite{Henk}, we find that every S-system is toric and can be described by:
\begin{equation}
\dot{X}_{i}(w_{1}, \ldots, w_{k}) =  
\alpha_{ip_{i}} \prod_{j=1}^{m} 
\left(w_{j}^{u_{j1}} w_{2}^{u_{j2}} \cdots w_{n}^{u_{jn}}\right)^{g_{ip_{i}}}
- 
\beta_{iq_{i}} \prod_{j=1}^{m}
\left(w_{j}^{u_{j1}} w_{2}^{u_{j2}} \cdots w_{n}^{u_{jn}}\right)^{h_{iq_{i}}}.
\end{equation}

Or, more compactly:
\begin{equation}
\dot{X}_{i}(w_{1}, \ldots, w_{k}) =  
\alpha_{ip_{i}} \prod_{j=1}^{m} (W_{i}^{h_{j}})^{g_{ip_{i}}}
-  
\beta_{iq_{i}} \prod_{j=1}^{m} (W_{i}^{h_{j}})^{h_{iq_{i}}}.
\end{equation}

We thus define a \textbf{Toric S-system} according to the expression above, where $h_{j}$ is an element of the Hilbert basis for some $j$.  
We assert that the binomials constructed from the Hilbert basis $H$,
\[
I_{H} = \langle W^{\alpha} - W^{\beta} \rangle,
\]
with $W_{i}^{h_{j} g_{ip_{i}}} = W^{\alpha}$ and $W_{i}^{h_{j} g_{iq_{i}}} = W^{\beta}$,  
are \textbf{toric binomials}. \textbf{q.e.d.}\\
\end{theorem}


  
%\begin{definition}\label{Definition 3.2.2} \textbf{(Toric variety)}. We define a \textbf{toric variety} associated to $ f $ as follows,\\
%\begin{equation}
%X_{\sigma} =Spec\left( R_{\sigma} \right) = \left\lbrace f_{1},....,f_{k}  \: \in \: R_{\sigma} \right\rbrace. \\
%\end{equation}\\

%where the notation for $ Spec\left( R_{\sigma} \right) $ means, the set of all the prime polynomials (non-divisible polynomials) $ \lbrace f_{1},....,f_{k} \rbrace $ in the ring $  R_{\sigma} $ for some integer $ k $. 

%\begin{definition}\label{Definition 3.2.3} \textbf{Environment cone.} Let $ f $ be an S-system of any rate law of a chemical concentration change of any species in a gene-metabolic network. We compute the support of $ f $ and the cone associated with it, $ \sigma \,\subseteq\,\mathbb{Z}^{n}$. We know that $ supp(f) \subseteq  \sigma $. We define an \textbf{environment cone} associated to $ f $ as the dual-cone $ \sigma^{\vee} $ of $ \sigma $, we also know that $supp(f) \subseteq  \sigma \subseteq  \sigma^{\vee} $.
%\end{definition}

%\textbf{Relevance in kinetic enzyme.} Up to here, we have equipped of new geometrical properties to the set of vectors associated to kinetic orders for an S-system, we have defined, cones, ring of polynomials, and a toric variety. But the most importance is the fact through of Gordan's lemma, see \cite{GEwald1993}. We have found a basis o generators (Hilbert basis) for all space of kinetic orders defined for any specie on the network. Now, we will try to exploit the properties gifted by the Hilbert basis.\\


%\textbf{Fix points on torus}. There exists several programms and algorithms to compute Hilbert basis, \cite{}, once obtained the set of generators $ H =\lbrace h_{1},..., h_{k} \rbrace $ for some cone $ \sigma =Con(b_{1},...,b_{r}) $ where each vector $ b_{j} $ is a face, \cite{},  that generates the cone. we define the fix points on the torus or toric variety through of the following formula, see \cite{}. \\

%\begin{equation}
 %X_{b_{j}} = \lim_{Z \longrightarrow 0} Z^{<b_{j},h_{i}>} =(\lim_{Z \longrightarrow 0}Z^{<b_{j},h_{1}>},...,\lim_{Z \longrightarrow 0}Z^{<b_{j},h_{k}>} ).
%\end{equation}

%We compute this limit for each face $ b_{j} $ in the cone, therefore, there are $ r $ fix points, or in another way, $  X_{b_{1}},..., X_{b_{r}} $. These limtis exists, if and only if, the set of constraints fullfil the conditions as we can see below.\\

%\begin{equation}
%<b_{j},h_{1}> >= 0,
%<b_{j},h_{2}> >= 0,
%...,
%<b_{j},h_{k}> >= 0.
%\end{equation}

%For some integers, $ j $ and $ k $, and usual scalar inner product, $ < , > $. We remember that the vectors $ b_{j} $ are of fact the \textbf{kinetic orders} associated to the support of some S-system $ f $ as we saw above. We use this set of constraints in addition with the constraints for dominant fluxes, to determine values for kinetic orders in the algorithm described in this work.\\

%\textbf{Assertion.} The classical point fixs, are information characterized by the well known steady-state from phenotype, in our case the fix points on torus, are information provided from genotype, we mean, this is another consequence of the monomial map, $g_{ip_{i}} \longmapsto X_{j}^{g_{ip_{i}}}    $ studied here. \\

%\textbf{Toric S-systems and dominant fluxes}.\\

%With the Hilbert basis at hand, $H =\lbrace h_{1},..., h_{k} \rbrace $, we give a matricial representation of it, if we consider each $ h_{i} $ as a row vector, thus we write, $h_{1} = (u_{11},u_{12},...,u_{1n}) $, $h_{2} = (u_{21},u_{22},...,u_{2n}) $,...,$h_{k} = (u_{k1},u_{k2},...,u_{kn}) $.\\\\

%\begin{center}
%$H  =   
%\begin{pmatrix}
%u_{11}&u_{12}&...&u_{1n}\\
%u_{21}&u_{22}&...&u_{2n}\\
%...&...&...&...\\
%u_{k1}&u_{k2}&...&u_{kn}
%\end{pmatrix}  \\\\
%$
%\end{center}\\\\


%To continuation, we can use the matrix defined above to represent the varibales $ X_{i} $ in a new system of coordinates, by means of the following monomial transformation, $ u_{ij} \longmapsto  w_{i}^{u_{ij}}$, see \cite{GEwald1993}, \cite{BSturmfels1996}, \cite{DCoxLittleSchenck2010}, the relevance to do that, is to notice dynamics properties encrypted for S-systems into of the Hilbert basis, \cite{BSturmfels1996}, \cite{Henk}. \\

%\begin{equation}  
%\begin{matrix}
%_{1}= w_{1}^{u_{11}}*&w^{u_{12}}_{2}*&...&*w^{u_{1n}}_{n},\\
%X_{2} =w_{1}^{u_{21}}*&w_{2}^{u_{21}}*&...&*w_{n}^{u_{2n}}\\
%...&...&...&...\\
%X_{k}=w_{1}^{u_{k1}}*&w_{2}^{u_{k2}}*&...&*w_{n}^{u_{kn}} \end{matrix}\\\\
%\end{equation}

%we can use this matrix representation above, to embedding to a new system of monomial coordinates for all family of S-system equations. Hence, we give the following result to extended the analysis of molecular networks towards a perspective to large-scale genetic and metabolic networks. \\


%\begin{theorem}\label{Theorem} \textbf{(Toric S-system)}. If we substitute this change of coordinates in the expression 2.2, the original S-systems, we have the following equation, it which is valid for all $ i=1,..,n $.\\

%\textbf{Proof:} Given an S-system representation for molecular networks, given by eq. (2.2). Using the Hironaka's theorem and the algorithmic process for Hilbert basis, a finite number of times over S-system equations, see \cite{MFAtiyah1970}, \cite{TFukui2000}, \cite{Henk}. Then, this process implies that all S-system is toric, is described by,

%\begin{equation}
%\dot{X}_{i}(w_{1},...,w_{k})=  \alpha_{ip_{i}} \prod_{j=1}^{m} (w_{j}^{u_{j1}}*w^{u_{j2}}_{2}*...*w^{u_{jn}}_{n})^{g_{ip_{i}}} -  \beta_{iq_{i}} \prod_{j=1}^{m}(w_{j}^{u_{j1}}*w^{u_{j2}}_{2}*...*w^{u_{jn}}_{n})^{h_{iq_{i}}}.\\
%\end{equation}
%Or simply, \\
%\begin{equation}
%\dot{X}_{i}(w_{1},...,w_{k})=  \alpha_{ip_{i}} \prod_{j=1}^{m} (W_{i}^{h_{j}})^{g_{ip_{i}}} -  \beta_{iq_{i}} \prod_{j=1}^{m}(W_{i}^{h_{j}})^{h_{iq_{i}}}.\\
%\end{equation}

%We define a Toric S-system, according to the expression given above, where $ h_{j} $ is an element of the Hilbert basis for some $j$, and we make the assertion that the binomials, built from the Hilbert basis (H),  $I_{H} = <W^{\alpha} - W^{\beta} >$ with $ W^{h_{j}*g_{ip_{i}}}_{i}= W^{\alpha} $ and $ W^{h_{j}*g_{iq_{i}}}_{i}= W^{\bet}$, they are toric binomials, \textbf{q.e.d}.\\
%\end{theorem}
The monomial transformation, \[ u_{ij} \longmapsto w_{i}^{u_{ij}}\] represents, in this geometric framework, the \textbf{Genotype–Phenotype relationship}. These are the coordinate transformations of the geometric basis corresponding to the kinetic orders such as the number of binding sites in regulatory genomic sequences, the order of biochemical reactions in metabolism, or the degree of multimeric regulation in proteins. 

The monomials $w_{i}^{u_{ij}}$ represent the chemical concentrations of the species or biomolecules described by the S-system $X_{i}$. The coordinates $u_{ij}$ encode information derived from the genome sequence, which is then expressed through the phenotypic gradient: \[\dot{X}_{j}(u_{ij}) = w_{i}^{u_{ij}}.\] Thus, the mapping of dynamical behavior occurs through this gradient, linking genotypic parameters to phenotypic observables.

% -------------------------------------------------------------------------
% Discussion Section
% -------------------------------------------------------------------------

\section{Discussion}

We have presented new topological properties of biochemical networks by exploiting the structural features of the Power Law and S-system modeling frameworks. In particular, we emphasize the significance of the monomial map: $g_{ij} \longmapsto X_{j}^{g_{ij}}$, which provides a bridge between biochemical kinetics and algebraic geometry. By focusing our analysis on the exponent space of S-systems, we study the kinetic orders using computational algebraic geometry techniques. This allows us to identify geometric bases that constrain and characterize the possible values of these parameters, thereby facilitating the solution of large systems of differential equations.

The central open questions that can be addressed with this approach are as follows:  
\begin{itemize}
    \item Under what conditions are these geometrical regions valid for assigning values to the kinetic orders?
    \item Which of these regions represent true biological information and which are merely mathematical artifacts?
\end{itemize}

Another advantage of the proposed methodology is its potential to model the dynamics of large-scale metabolic and gene-regulatory networks. The monomial map has a formal justification through the mathematical concept of a \textbf{toric variety}. Although toric varieties are abstract algebraic objects, they serve as powerful tools for modular analysis: each variety corresponds to a submodule of the biochemical network, and the full network can be reconstructed by \textit{gluing together} these varieties using geometric techniques.

While a single toric variety may provide limited insight into the overall biochemical system, the ensemble of all varieties properly joined captures the global dynamics of large-scale genetic and metabolic networks. This modular, geometrically grounded framework thus offers a new perspective for understanding and simulating the complex interplay between genotype and phenotype.

Finally, we have included in the Supplementary Information the complete set of equations used for the toric analysis of a gene–metabolic network in the allosteric system of the tetrameric protein TrpR (tryptophan repressor) in E. coli K-12. In a future work, we will solve the complete dynamics using toric varieties such as biochemical modules.


%The monomial transformation $  u_{ij} \longmapsto w_{i}^{u_{ij}} $ represents in this geometry, the relatioship \textbf{Genotype-Phenotype}, they are coordinates of geometrical basis for the kinetic orders (number of binding sites in regulatory genome sequences, order of biochemical reactions in metabolism or multimeric regulation sub units in proteins), the monomials $ w_{i}^{u_{ij}} $ are chemical concentrations of any species or biomolecules described by the S-system $X_{i}  $, the coordinates $ u_{ij}  $ are information from genome sequence it which is read by the phenotypic gradient $ \.{X}_{j}(u_{ij})=w_{i}^{u_{ij}} $.
%as we saw in the different scenaries in the fig. 4, the dynamcis are mapped by this gradient.\\

%\textbf{Dominant Fluxes in toric S-systems.} Now we consider the equation (2.1), and we use the monomial transformation (2.9), we substitute the new coordinates system in (2.1),\\

%\begin{equation}
%\dot{X}_{i}= \sum_{k=1}^{p_{i}} \alpha_{ik} \prod_{j=1}^{m} (W_{i}^{h_{j}})^{g_{ijk}} - \sum_{k=1}^{q_{i}} \beta_{ik}\prod_{j=1}^{m} (W_{i}^{h_{j}})^{h_{ijk}},\: i=1,...,n.
%\end{equation}

%We run some terms on the sum, thus we have the following formal expression,\\
%\begin{equation}
%\dot{X}_{i}= \alpha_{i1} \prod_{j=1}^{m} (W_{i}^{h_{j}})^{g_{ij1}} +\alpha_{i2} \prod_{j=1}^{m} (W_{i}^{h_{j}})^{g_{ij2}} + \alpha_{i3} \prod_{j=1}^{m} (W_{i}^{h_{j}})^{g_{ij3}}+.... - \\\\
%\beta_{i1}\prod_{j=1}^{m} (W_{i}^{h_{j}})^{h_{ij1}}-

%beta_{i2}\prod_{j=1}^{m} (W_{i}^{h_{j}})^{h_{ij2}}-\beta_{i3}\prod_{j=1}^{m} (W_{i}^{h_{j}})^{h_{ij3}}-.....,
%\end{equation}

%We get several inner products in the exponents of this expression, as we show below,\\

%$ \left\langle h_{j} , g_{ij1}\right\rangle $, $ \left\langle h_{j}, g_{ij2}\right\rangle $, $ \left\langle h_{j}, g_{ij3}\right\rangle...$, $ \left\langle h_{j},h_{ij1} \right\rangle  $, $ \left\langle h_{j},h_{ij2} \right\rangle  $, $ \left\langle h_{j} ,h_{ij3} \right\rangle.... $. \\


%\section{Discussion.}

%We have shown new topological properties in biochemical networks, exploting the characteristics of Power Law and S-systems modeling, we notice mainly the monomial map property, $ g_{ij} \longrightarrow X^{gij}_{j} $. we take advantage of this map analysing only the exponent space in S-systems, as consequence we focus our analysis to kinetic orders by means of computational algebraic geometry techniques to find geometrical basis to bound the values of these parameters. This technique let us to solve big set of differential equations. The new issues to solve with this technique, are : when are valid these regions to give values to kinetic orders?, what geometrical regions are mathematical artifacts and which are regions with biological information for the netwok analysis.\\
%Another of the advantages of the methodology presented here, is the possibilty of modeling for dynamics in large-scale networks in metabolism and gene circuitry. The property of monomial map has the formal justyfication to module netowrks through the mathemtical concept of \textbf{toric variety}, they are abstract mathematical objects, but they are capable to give information in modules, to analyse module by module through varieties and after to glue thogether in a big toric variety. Pherphaps, the information that provides one variety is not very informative for the biochemical network, but the whole pieces or varierties glueded togheter let us see the complete dynamics in large-scale networks. 

 

\section{Acknowledgment}
Marco Polo Castillo Villalba is a doctoral student from the Programa de Doctorado en Ciencias Biomédicas, Universidad Nacional Autónoma de México (UNAM) and has received the CONAHCYT fellowship 754050. We acknowledge funding from UNAM and from the National Institutes of Health (grant number 5R01GM110597-03). I give a special thanks to Michael A. Savageau for all technical comments during my scholar stay with him, Pedro Miramontes Vidal and Julio Collado-Vides and technical contribution by Laura Gómez Romero.

%%%%%%%%%%%%%%%%%%%%%%%%%%%%%%%%%%%%%%%%%%%%%%%%%%%%%%%%%%%%%
\section*{Declaration of Interest Statement}

The author declares that he has no personal, financial, professional, or 
academic conflicts of interest.
%%%%%%%%%%%%%%%%%%%%%%%%%%%%%%%%%%%%%%%%%%%%%%%%%%%%%%

\begin{thebibliography}{1}

\bibitem{MFAtiyah1970}
M.F.~Atiyah.
%\newblock Information theory and an extension of the maximum likelihood
 % principle.
{\em Resolution of Singularities and Division of Distributions}, Communications On Pure and Applied Mathematics, vol. XXIII, 145-150 (1970).

\bibitem{DCoxLittleOshea2004}
D.~Cox, J.~Little and D.~O'Shea.
%\newblock Information theory and an extension of the maximum likelihood
 % principle.
{\em Using Algebraic Geometry.} Springer, 2nd. Edition, August, 2004.


\bibitem{DCoxLittleSchenck2010}
D.~Cox, J.~Little and H.~Schenck.
%\newblock Information theory and an extension of the maximum likelihood
 % principle.
{\em Toric Varieties} Department Of Mathematics, Amherst College, Amherst MA 01002, 2010.

\bibitem{GEwald1993}
G.~Ewald.
%\newblock Combinatorial Convexity and Algebraic Geometry.
%\newblock In T.M. Fomby and R.~Carter Hill, editors, 
{\em Combinatorial Convexity and Algebraic Geometry.}, Springer-Verlag, New York, Inc., 1996.

\bibitem{Henk} 
Henk M, Weismantel R. 
\newblock On Hilbert bases of polyhedral cones.
{\em Konrad-Zuse-Zentrum für Informationstechnik; 2010.}

\bibitem{TFukui2000}
T.~Fukui.
\newblock Introduction to Toric Modifications With an Application to Real Singularities.
{\em Department Of Mathematics, Faculty Of Science, Saitama University}. 255 Shimo-Okubo, Urawa 338-8570, Japan, pag. 96-114, 2000.

\bibitem{LomnitzSavageau2013}
J. G.~Lomnitz, M. A.~Savageau.
\newblock Phenotypic deconstruction of gene circuitry
%\newblock In T.M. Fomby and R.~Carter Hill, editors, 
{\em Chaos.} Vol. 23, Issue 2, May 21, 2013.
\bibitem{LomnitzSavageau2014}
J. G.~Lomnitz, M. A.~Savageau.
\newblock Strategy Revealing Phenotypic Differences among Synthetic
Oscillator Designs.
%\newblock In T.M. Fomby and R.~Carter Hill, editors, 
{\em ACS Synth. Biol.} Volume 3, 686−701, July 14, 2014.

\bibitem{LomnitzSavageau2015}
J. G.~Lomnitz, M. A.~Savageau.
\newblock Elucidating the genotype-phenotype map by automatic
enumeration and analysis of the phenotypic repertoire.
%\newblock In T.M. Fomby and R.~Carter Hill, editors, 
{\em npj Systems Biology and Applications.} Vol 1, 15003; doi:10.1038/npjsba.2015.3; published online 28 September 2015.
\bibitem{LomnitzSavageau2016}
J. G.~Lomnitz, M. A.~Savageau.
\newblock Design Space Toolbox V2: Automated
Software Enabling a Novel
Phenotype-Centric Modeling
Strategy for Natural and Synthetic
Biological Systems.
%\newblock In T.M. Fomby and R.~Carter Hill, editors, 
{\em Frontiers in Genetics.} Vol 27, article 117, 12 July, 2015.

\bibitem{MASavageau1976}
%\input amsart-template.bbl
\newblock Biochemical Systems Analysis: “A Study of Function and Design in Molecular Biology”.
%\newblock In T.M. Fomby and R.~Carter Hill, editors, 
{\em Biochemical Systems Analysis.}, volume 1. Addison-Wesley Inc , Reading, Mass, USA, 1976.

\bibitem{SavageauOptimal} 
Savageau MA. 
\newblock Optimal design of feedback control by inhibition. Dynamic considerations. 
{\em J Mol Evol.} 1975;5:199–222.

\bibitem{MASavageauVoit1987}
M. A.~Savageau and E. O.~Voit.
\newblock Recasting Nonlinear Differential Equations as S-Systems:
A Canonical Nonlinear Form.
%\newblock In T.M. Fomby and R.~Carter Hill, editors, 
{\em Mathematical Biosciences.}, Volume 87, Issue 1, pages 83-115, November 1987.
\bibitem{MASavageauVoitIrvine987}
M. A.~Savageau, H. O.~Voit and D. H.~Irvine.
\newblock Biochemical Systems Theory and Metabolic Control Theory:
1. Fundamental Similarities and Differences.
%\newblock In T.M. Fomby and R.~Carter Hill, editors, 
{\em Mathematical Biosciences.}, Volume 86, Issue(2), pages 127-145, October 1987. 
\bibitem{MASavageau2001}
M. A.~Savageau.
\newblock Design principles for elementary gene circuits: Elements, methods, and examples.
%\newblock In T.M. Fomby and R.~Carter Hill, editors, 
{\em Chaos.}, Volume 11, Issue 1, December (2001).

\bibitem{SavageauCoelhoFasaniSalvador2009}
M. A.~Savageau, P. M. B. M.~Coelho, R. A.~Fasani  and A.~Salvador.
\newblock Phenotypes	and	tolerances	in	the	design	space
of	biochemical	systems.
%\newblock In T.M. Fomby and R.~Carter Hill, editors, 
{\em PNAS.} Vol. 106, no. 16, 6435– 6440, April 21, 2009.

\bibitem{BSturmfels1996}
B.~Sturmfels.
\newblock Grobner Bases and Convex Polytopes.
%\newblock In T.M. Fomby and R.~Carter Hill, editors, 
{\em American Mathematical Society.} Providence, RI, 1996.

\bibitem{rev15}
Voit. E. O.
Non Linear Canonical Modeling. S-system Approach To Understanding Complexity.
New York. New Nonstrand Reinold, (1991).

\end{thebibliography}

\section*{Support information}

\begin{figure}[h!]
\centering
\makebox{\includegraphics[height=7.25in]{TrpR.png}}
\caption{Tryptophan genetic and metabolic network}
\end{figure}

\begin{figure}[h!]
\centering
\makebox{\includegraphics[height=3.25in]{Toric_analysis1.png}}
%\caption{Toric S-system equations}
\end{figure}

\begin{figure}[h!]
\centering
\makebox{\includegraphics[height=2.25in]{Toric_analysis2.png}}
\caption{Toric S-system equations for tetrameric tryptophan protein.}
\end{figure}


%\input amsart_template.bbl
\end{document}
