## Supplementary figures and images for "Towards Modeling in Large-Scale Genetic and Metabolic Networks"

### plot_paper1.gif

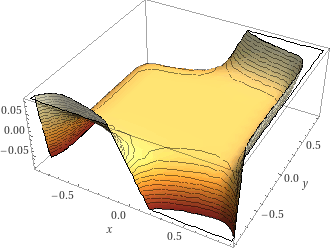

### plot_paper1.jpg

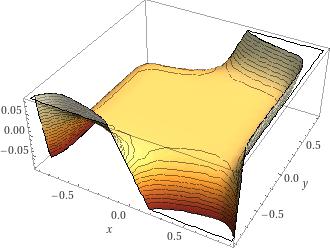

### plot_paper2.gif

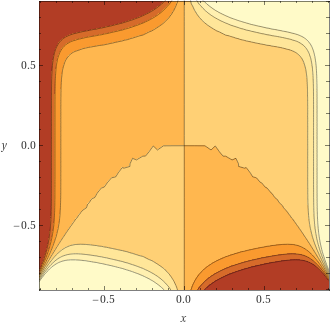

### plot_paper2.jpg

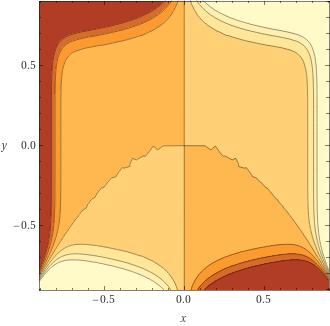

### plot_paper3.gif

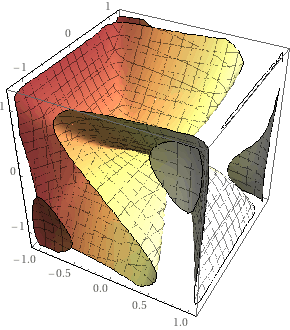

### plot_paper3.jpg

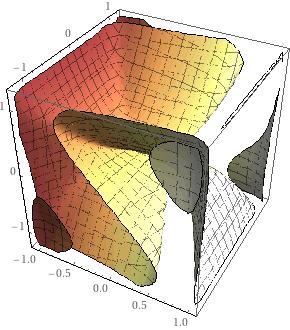

### plot_paper4.gif

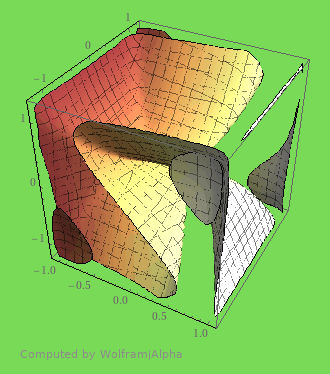

### plot_paper4.jpg

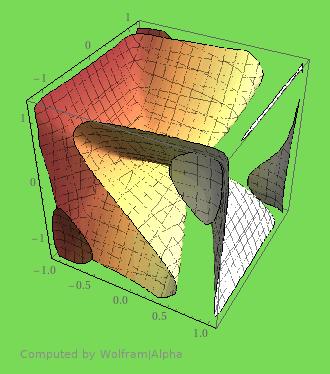

### plot_paper5.gif

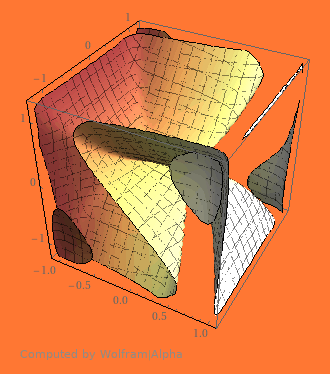

### plot_paper6.gif

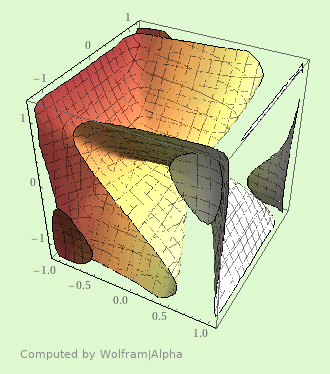

### Proteins_paper.jpg

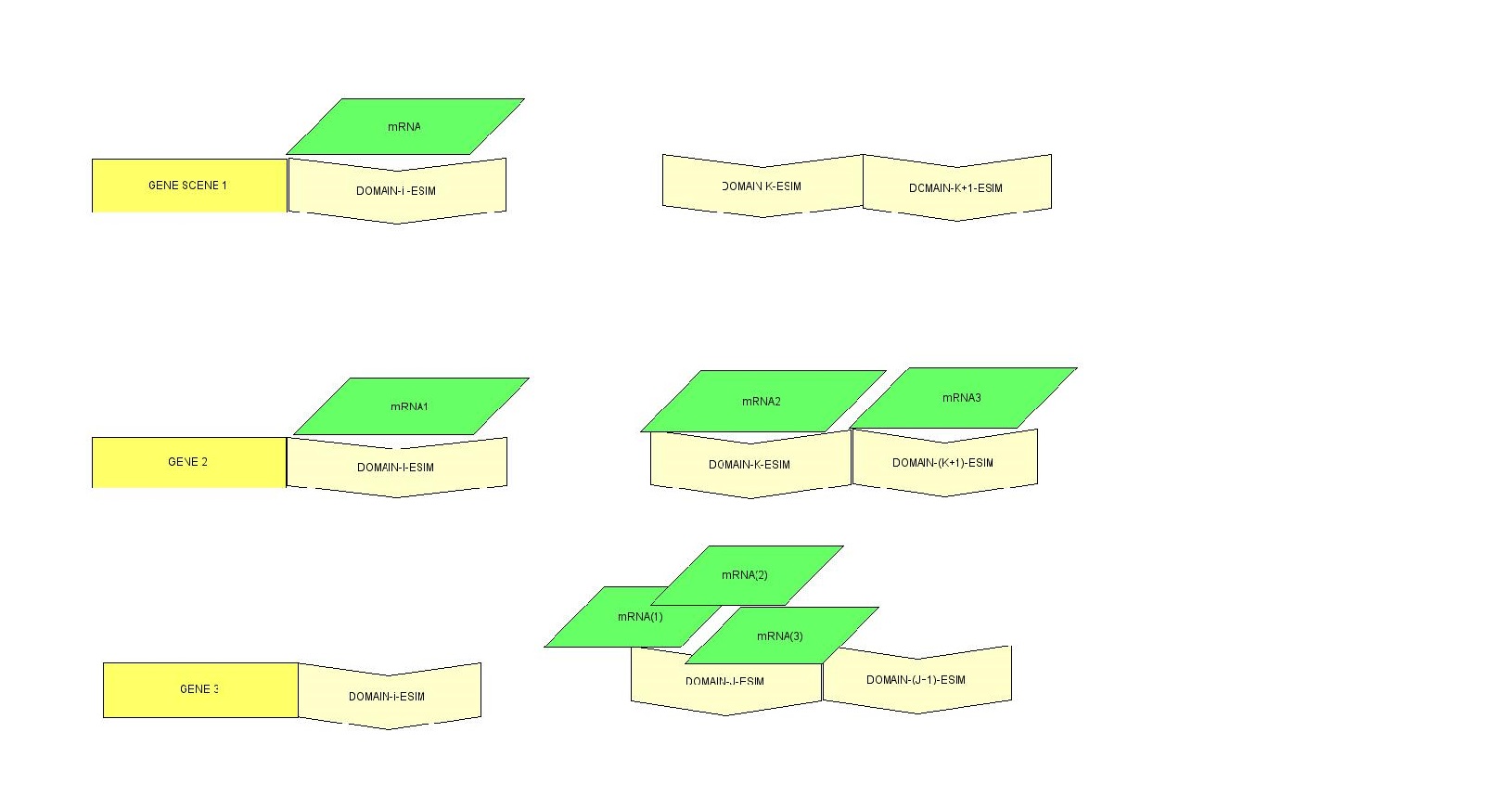

### Proteins_sceneries.png

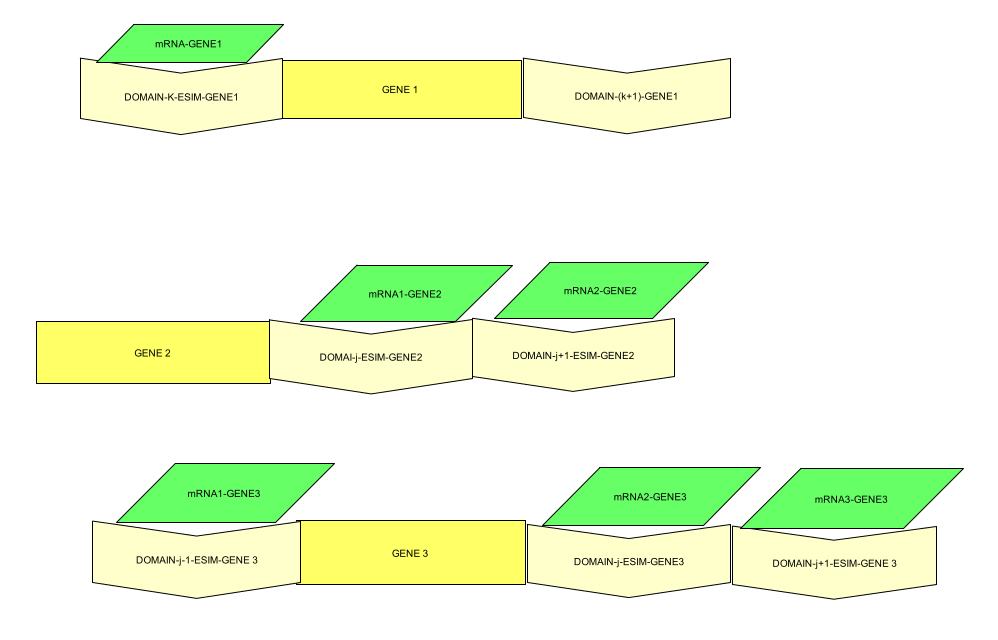

### region1.png

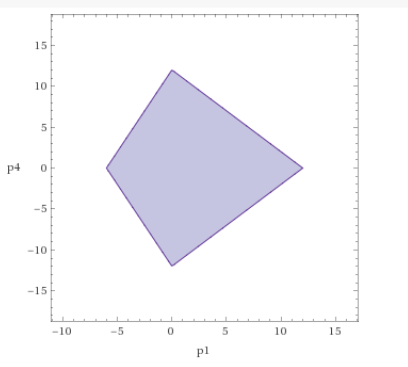

### region2.png

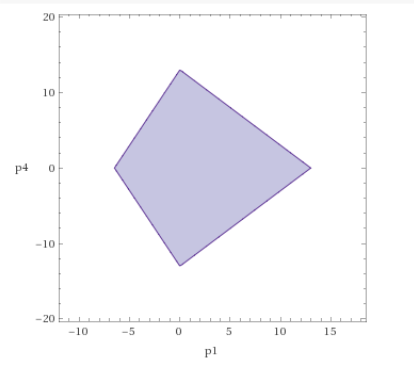

### Toric_analysis1.png

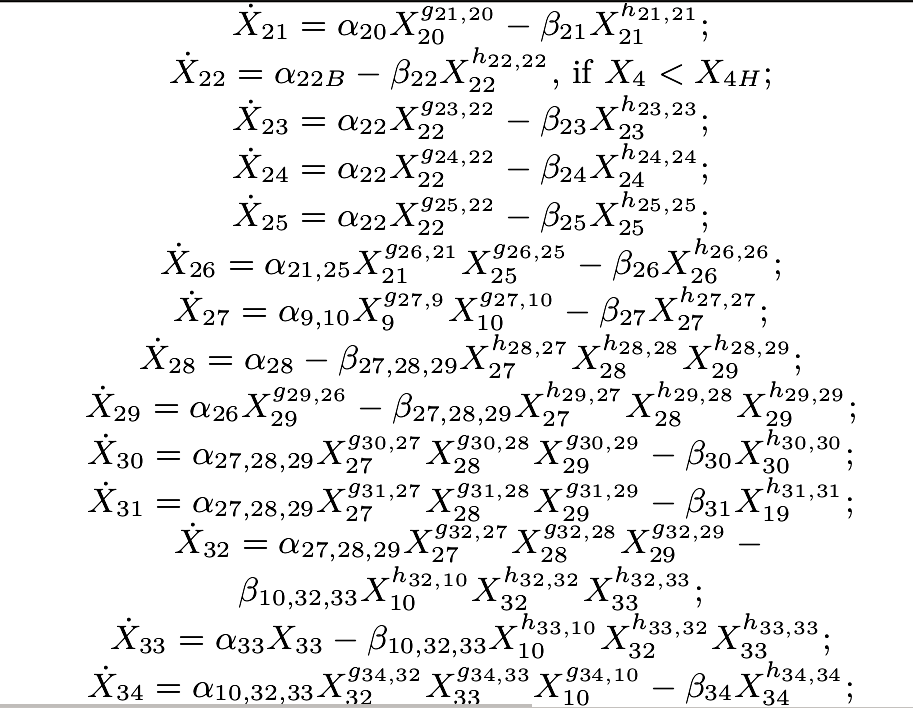

### Toric_analysis2.png

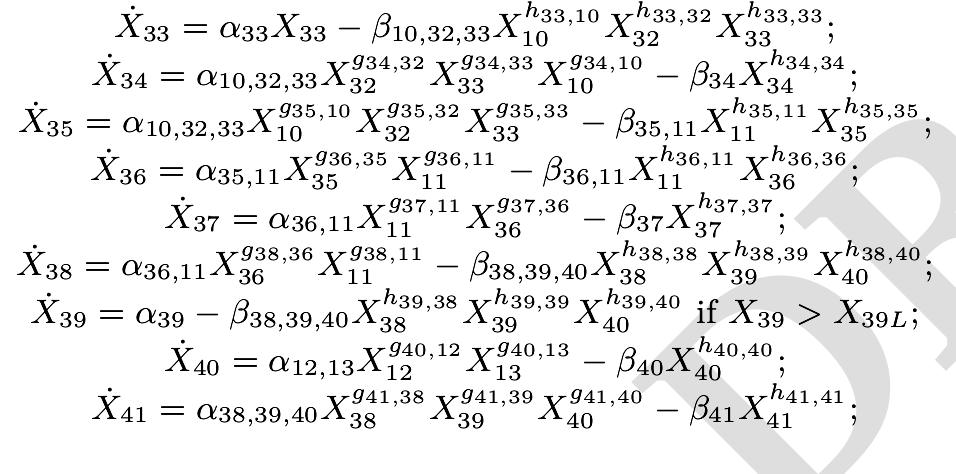

### toy_circuit.png

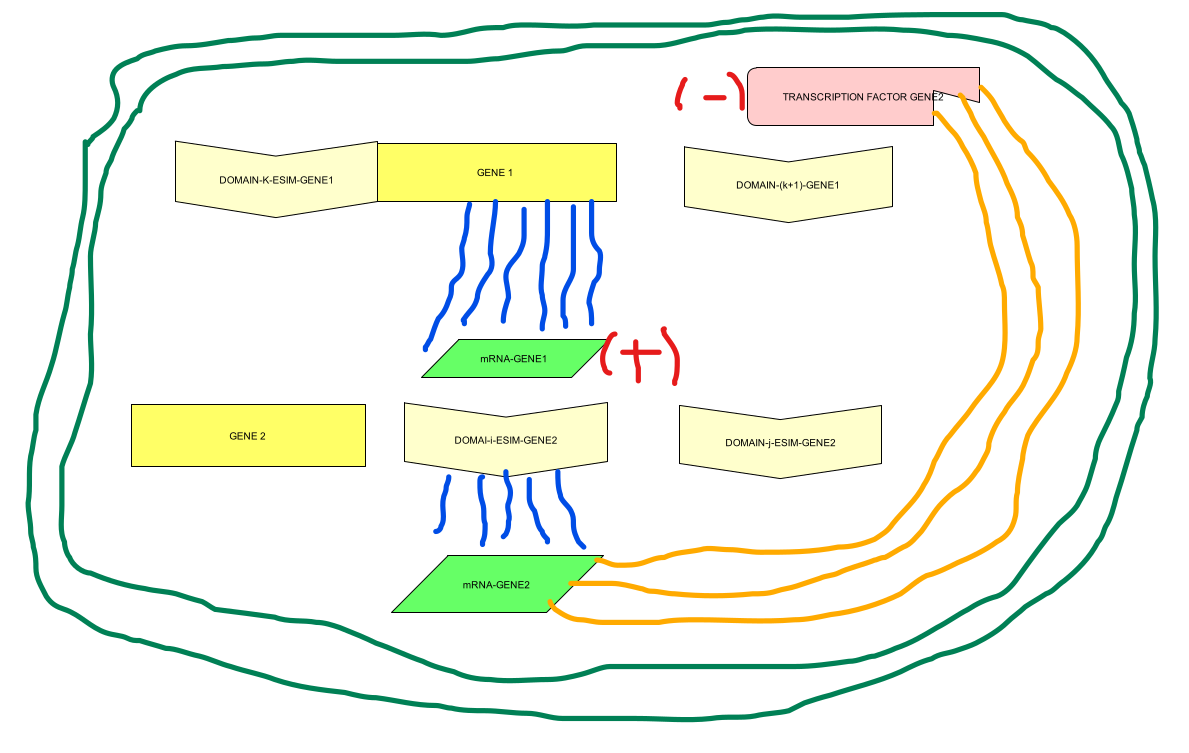

### TrpR.png

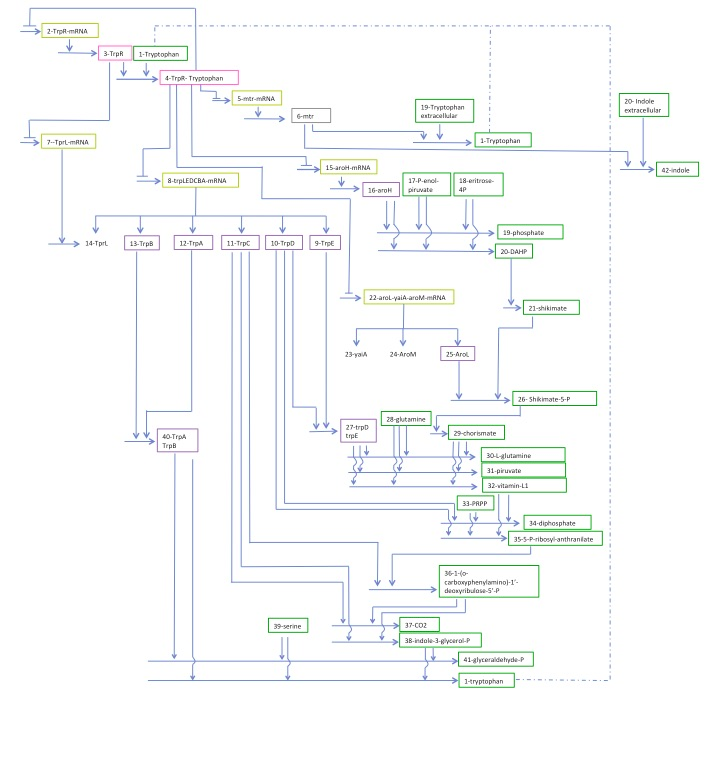
